# EMT and cell cycle control invadopodia and metastasis in breast cancer via Filip1L

**DOI:** 10.64898/2025.12.05.692654

**Authors:** Elizaveta Belova, Afrooz Jarrah, Gleb Abalakov, Adam L Karami, Bojana Gligorijevic

## Abstract

Invadopodia are actin- and protease-rich membrane structures that enable cancer cells to degrade extracellular matrix (ECM). Invadopodia activity is cell cycle-dependent, but how their regulation across the cell cycle is influenced by epithelial-to-mesenchymal transition (EMT) remains unclear. We show that as the EMT progresses, cell engagement in invadopodia-mediated ECM degradation shits from G2 phase in Early E/M cells to G1 phase in Late E/M or M cells.

Using bulk mRNA sequencing of Early- versus Late- E/M cells sorted by cell cycle phase, we identified FILIP1L as an EMT- and cell cycle-regulated candidate whose expression peaks in the invasive phase of each cell state: G2 in Early E/M cells and G1 in Late E/M cells.

We next demonstrated that FILIP1L is a novel invadopodia component, whose loss increases ECM degradation while impairing migration and 3D invasion. In mouse models, FILIP1L KD tumors develop fewer metastatic colonies, suggesting that FILIP1L supports productive invasion by coordinating between invadopodia and migratory cell states. FILIP1L expression increases with EMT progression and correlates with poor outcomes in breast cancer patients. Together, these findings identify a previously unrecognized link between EMT, cell cycle and invadopodia and establish FILIP1L as the key regulator of this process.

## Introduction

Breast cancer remains one of the most common malignancies among women, with an overall 5-year survival rate of 87–99% for localized and regional stages. However, once the disease progresses to the metastatic stage with distant metastases, the 5-year survival rate drops significantly to about 32% [1]. Current treatment options for metastatic breast cancer are very limited and primarily palliative, aiming to control symptoms and slow disease progression rather than achieve a cure. This highlights the critical need to develop new therapeutic strategies that can effectively limit metastasis and extend patient survival.

Distant metastasis is the leading cause of breast cancer-related deaths. Despite the availability of numerous treatments, such as surgery and chemotherapy, and immunotherapeutic approaches, approximately one-third of patients ultimately develop metastatic disease [2]. Primary sites of breast cancer metastasis are bone, lungs, brain, and liver [3]. Some patients also develop metastasis in multiple sites [4].

To successfully disseminate to distant organs, breast cancer cells undergo a number of sequential steps in the metastatic cascade. This includes local invasion into the stromal extracellular matrix (ECM) rich in collagen I, breaching the basement membrane which surrounds the blood vessels, intravasation into the bloodstream, circulation in the blood flow, extravasation into the distant organ and growth of metastatic colonies. In order to intravasate and extravasate, metastatic cells require a actin-rich membrane protrusions with the proteolytic ability referred to as invadopodia [5–8].

In recent years *in vivo*, *in vitro* and clinical studies of epithelium-derived malignancies, including breast carcinomas, have pointed out to the importance of the epithelial-mesenchymal transition (EMT) during tumor progression and the high potential of the hybrid epithelial/mesenchymal (E/M) states for the cancer cells dissemination. EMT is a developmental program during which the tightly packed epithelium transitions into loosely connected mesenchymal state [9]. In cancer, EMT is usually associated with increased invasiveness, chemoresistance and poor prognosis [9–14]. One of the reasons for this is the increased ability of EMT-undergoing cells to assemble invadopodia [15,16] and degrade ECM [17–19]. The EMT process is more accurately described as a spectrum of states. The reverse process, known as mesenchymal-to-epithelial transition (MET), can also occur, often when cells have reached secondary organs [20]. While most studies focused on extremes of the EMT transition, some cells are stable in partial a.k.a. hybrid E/M states along the EMT spectrum, which is characterized by the presence of the intercellular adhesions and the simultaneous expression of epithelial (E) state markers such as E-cadherin, ZO1, cytokeratin, as well as mesenchymal (M) state markers such as vimentin, fibronectin, and expression of EMT-associated transcription factors, including Snail, Twist and Zeb families [20–22]. Our recent work showed that during the collective 3D spheroid invasion, hybrid E/M cells assemble invadopodia in the leader cells, which are located at the tip of the invading strand. When mixed with invadopodia-deficient cells, E/M leader cells allow for cooperative invasion and metastasis [16].

The 3D invasion is a complex process that consists of oscillations between two distinct motility-related states: invadopodia state, during which cells assemble invadopodia and degrade ECM, while remaining relatively stationary, and migratory state, during which cells translocate through the degraded ECM, using contractility machinery [23]. In the two-dimensional (2D) culture assays, such as the gelatin degradation assay, the two states are observed as mutually exclusive in time and space, and the elimination of invadopodia state still allows for cell to migrate laterally across the dish [24]. In this model, elimination of migration state results in stationary cells with long-lasting, highly degradative invadopodia structures [24,25]. However, in the more physiologically relevant 3D environments such as the 3D spheroid invasion, where cells are embedded in dense collagen I [26,27] invasion is tightly integrated, and requires the presence of both invadopodia- and migratory- states. Disruption of either state leads to impair of overall invasive capacity in 3D [23,24,28].

Both invadopodia and 3D invasion are affected by a number of extrinsic and intrinsic factors [29],one of which is cell cycle progression. We and others showed that in the mesenchymal cancer cells, p27 controls invadopodia-mediated degradation. As a result, ECM degradation occurs predominantly during the G1 phase of the cell cycle [25,30,31]. To maintain ECM degradation while progressing through cell cycle, leader cells in mesenchymal 3D spheroids forfeit their position when they transition from G1 to S phase, a process called leader-follower transition [25,32].

Invadopodia turnover rate greatly influences the extent of ECM degradation. In 2D culture systems, reduced invadopodia turnover results in fewer but larger degradation areas, because of prolonged matrix engagement and more stable structures [25,33]. While this may enhance localized degradation, it also reduces the spatial distribution of proteolytic activity across the substrate. In the context of 3D invasion, such decreased turnover impairs the cell’s ability to adaptively remodel the surrounding ECM in a spatially coordinated manner and results in lowered spheroid outgrowth and invasive progression [23].

In this study, we aim to investigate whether the coupling between cell cycle and invadopodia-mediated invasion is conserved across EMT states. As a model for E/M cells, we utilized 4T1 cell line. Surprisingly, our data demonstrates that 4T1 E/M cells degrade the ECM and invade in 3D during the G2 phase of the cell cycle. We have established that the EMT-inducing TGFβ treatment transitions E/M 4T1 cells to a later E/M state, characterized by increase in aspect ratio but preserving E-cadherin expression. We also induced the overexpression of EMT-related Snail 2 i.e. Slug transcription factor in 4T1 cells, leading to a full transition to M state. In Late E/M or M cells, ECM degradation and invasion shift from occurring primarily during the G2, to occurring in G1 phase of the cell cycle. Using cell cycle phase-specific RNASeq, we demonstrate that the shift from G2- to G1- degradation is driven by the cell cycle-dependent expression of a novel invadopodia component, Filamin A Interacting protein 1-like (FILIP1L). Knockdown of FILIP1L increases invadopodia-driven ECM degradation, while reducing cell migration, 3D spheroid invasion, and the number of metastatic lung colonies in a preclinical mouse model. *In silico* analysis of TCGA database show that increased FILIP1L expression correlates with shorter disease-free and progression-free survival outcomes in breast cancer patients.

In summary, we have identified an EMT- and cell cycle-sensitive component of invadopodia, FILIP1L, linking EMT progression to cell-cycle dependent control of invasive behavior. Given the essential role of invadopodia in intravasation and, consequently, in breast cancer metastasis, inhibitors of FILIP1L may represent a promising therapeutic approach when used in combination with conventional anti-proliferative chemotherapies.

## Results

### Invadopodia-mediated ECM degradation occurs during G2 phase in Early E/M cancer cells, and during G1 phase in Late-E/M and M cells

To enable real-time visualization of different phases of the cell cycle for studies of the cell cycle regulation of invadopodia in E/M cells, we generated 4T1-FUCCI cell line, which stably expresses the nuclear FUCCI reporters. Immunolabeling for invadopodia marker Tks5 in 4T1-FUCCI confirmed colocalization of Tks5 with gelatin degradation sites, confirming that 4T1-FUCCI cells actively degrade the ECM while in the G2 phase (**Figure 1A**).

**Figure 1.**
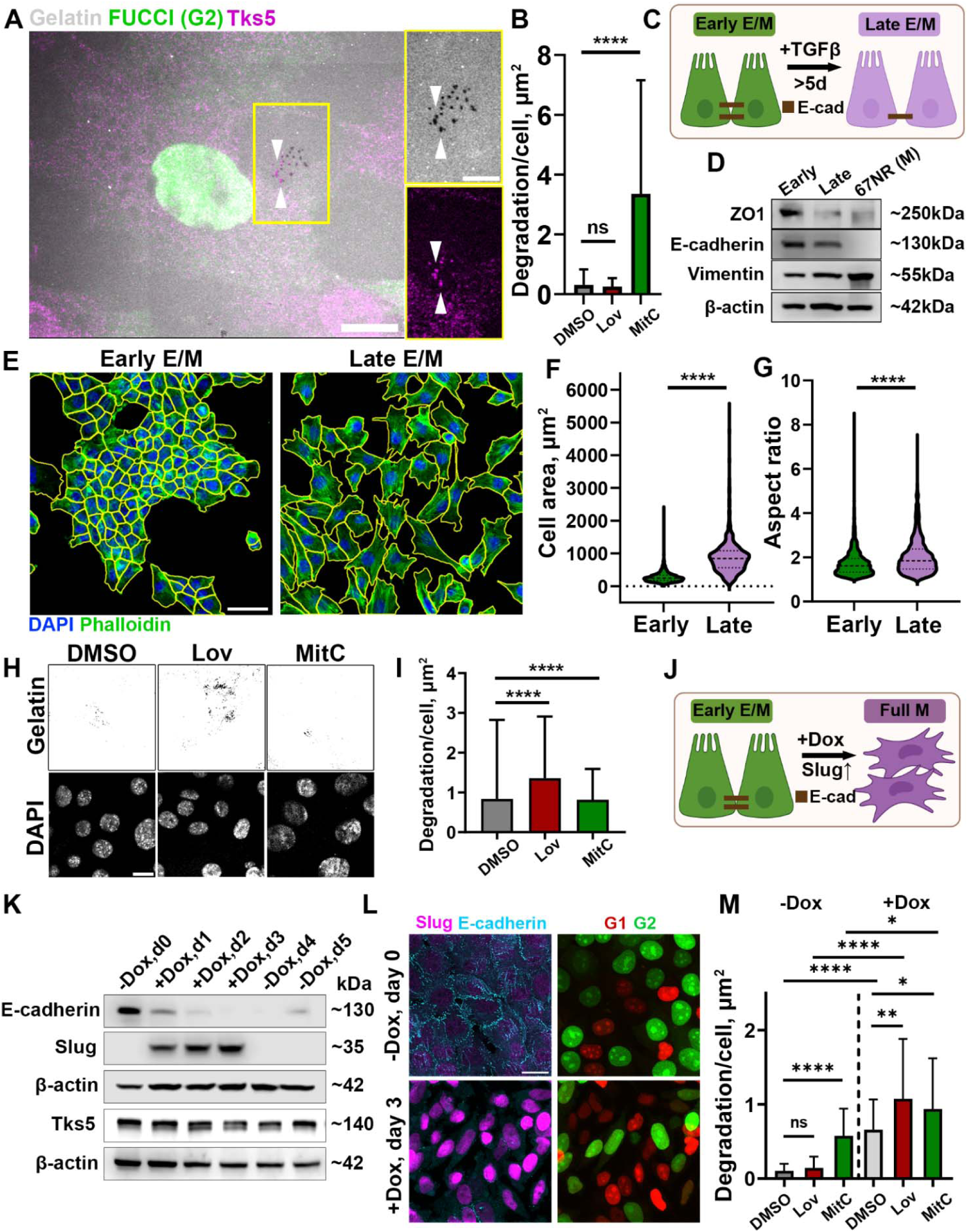
In early E/M cells, invadopodia degrades ECM during G2 phase, while in late E/M degradation occurs during G1 phase of the cell cycle. **A.** Early E/M 4T1-FUCCI cells synchronized in G2 using mitomycin C (nuclear green FUCCI) were cultured on fluorescent gelatin (gray) and immunolabeled for Tks5 (magenta). Scale bar: 20 µm. The inserts show a zoom-in of the boxed area, while arrowheads indicate representative puncta where degradation (top) colocalizes with invadopodia (bottom, Tks5). Scale bar: 10□µm. **B.** Invadopodia degradation in Early E/M cells: unsynchronized (DMSO) or synchronized in G1 by lovastatin (Lov), or in G2 phase (MitC). Bar height represents the mean, and error bars represent SD. N=3, n=1-3, 3-11 FOVs per n. **C.** Schematics of Late E/M induction via TGFβ-1 treatment. Created in BioRender. **D.** Western blot of epithelial markers (ZO1, E-cadherin) and mesenchymal marker (Vimentin) of Early E/M and Late E/M (TGFβ-1-treated) states. Mesenchymal (M) 67NR cells were used as a control. **E.** Morphological features of Early vs Late E/M states. Cells were labeled with DAPI (nuclei, blue) and phalloidin (cell borders, green). Yellow outlines represent results of Cellpose segmentation. Scale bar: 100 µm. **F.** Measurements of cellular areas in Early vs Late E/M cells. ****P <0.0001, Mann–Whitney U test. Dotted lines represent quartiles; the dashed line represents the median. N=3, n=3, 768 cells total. **G.** Aspect ratios in Early vs Late E/M cells. ****P <0.0001, Mann–Whitney U test. Dotted lines represent quartiles; the dashed line represents the median. N=3, n=3, 768 cells total. **H.** Representative images of gelatin degradation of Late E/M cells: unsynchronized (DMSO) or synchronized in G1 by lovastatin (Lov), or in G2 phase (MitC). **I.** Quantification of invadopodia degradation in Late E/M cells. Bar height represents the mean, and error bars represent SD. N=4, n=1-3, 5-24 FOVs per n. **J.** Schematics of full EMT induction via Slug overexpression. Created in BioRender. **K.** Western blot analysis of E-cadherin, Slug, and Tks5 in 4T1-FUCCI-TetOn-Slug cells. **L.** E-cadherin (cyan) and Slug (magenta) in 4T1-FUCCI-TetOn-Slug cells, -Dox and +Dox, (left panels and FUCCI G1 (Red), FUCCI G2 (Green) (right panels). Scale bar: 20 µm. **M.** Invadopodia degradation in 4T1-FUCCI-TetOn-Slug cells, -Dox- and +Dox, either unsynchronized (DMSO), synchronized in G1 (Lov), or G2 phase (MitC). Bar height represents the mean, and error bars represent SD. N=3, n=1-3, 5-10 FOVs per n.

We synchronized cells in G1 phase using lovastatin or in G2 phase, using mitomycin C [25]. Unexpectedly, cells synchronized in G1 were approximately 4-fold less efficient in ECM degradation compared to the unsynchronized cells, and 20-fold less efficient compared to those synchronized in G2 (**Figure 1B)**. These results suggest that, unlike MDA-MB-231 and BT-549 mesenchymal (M) cells [25], hybrid E/M cells degrade ECM mainly during the G2 phase of the cell cycle.

We hypothesized that the EMT status might be the causing factor of this opposing trend in cell cycle control of invasion. To test this, we employed a 5+ day TGFβ-1 treatment (**Figure 1C**), an established treatment to induce EMT through activation of the EMT-related transcription factors, towards a more mesenchymal state in 4T1 cells [34,35].

Cells treated by TGFβ-1 showed an increase in the cellular area (**Figure 1E, F**) and aspect ratio (**Figure 1G**), changes consistent with EMT transition from a rounded epithelial shape to a spindle-like, mesenchymal morphology. However, Western blot analysis showed that while the TGFβ-1 treatment led to the downregulation of the epithelial markers ZO1 and E-cadherin, a low-level of E-cadherin expression was maintained (**Figure 1D**). Hence, we termed untreated cells as cells in Early E/M state and TGFβ-1-treated cells as cells in Late E/M state.

Next, we tested if the transition to the Late E/M state led to a change in cell cycle regulation of invadopodia-mediated ECM degradation. Consistent with a shift from Early- to Late- E/M state, synchronized Late E/M cells demonstrated that the ECM degradation was performed predominantly during G1-state, with a minor portion of degradation being performed during G2 -state (**Figure 1F, G**). We wondered whether a full EMT, characterized by a complete E-cadherin downregulation, would result in ECM degradation being done solely in G1. To test this, we introduced a stable, doxycycline-inducible Slug overexpression into 4T1 cells (4T1-tetO-Slug) [36]. Slug is a transcription factor upregulated in BRCA1-mutated breast tumors [37] and known to promote metastasis [38,39]. Upon induction of Slug overexpression with 2 μM doxycycline (Dox), E-cadherin downregulation was observed on day 3 (**Figure 1H**). Withdrawal of Dox from the culture medium led to a loss of Slug expression on the following day (-Dox, d4), with E-cadherin expression beginning to recover by day 2 post Dox withdrawal (-Dox, d5) (**Figure 1H**). When -Dox and +Dox cells were synchronized in G1- or G2 - state, the highest degradation was observed in +Dox cells synchronized in G1 (lovastatin), corroborating findings from the TGFβ-1 treated cells (**Figure 1J, Figure S1A**). Additionally, Slug overexpression resulted in an increased proportion of cells in the G1 phase (**Figure S1B, C**). Interestingly, unsynchronized +Dox cells exhibited higher overall degradation compared to unsynchronized -Dox cells (-Dox DMSO vs. +Dox DMSO) even though Slug overexpression did not alter Tks5 levels (**Figure 1H**).

To determine whether TGFβ-1 treatment promotes an epithelial–mesenchymal (E/M) phenotype in epithelial cells (**Figure S2A**), we treated normal human mammary epithelial MCF10A cells with TGFβ-1 for 5 days and then plated them on gelatin matrix. This cell line was previously shown to assemble invadopodia and degrade ECM upon treatment with TGFβ-1 [40]. Compared to untreated cells, TGFβ-1-treated MCF10A cells exhibited elongated morphology and increased punctate gelatin degradation (**Figure S2B**). Quantification of the degradation area per cell revealed a significant increase in gelatin degradation upon TGFβ-1 treatment compared to the untreated cells (**Figure S2C**). When TGFβ-1-treated cells were synchronized in G1 phase using lovastatin, the degradation activity was markedly reduced, aligning with the results obtained for Early E/M 4T1 cells. To confirm the induction of EMT, we analyzed epithelial and mesenchymal marker expression by western blotting. Treatment by TGFβ-1 did not affect E-cadherin expression but has increased vimentin levels compared to the untreated cells, confirming a shift from E to an Early- E/M phenotype (**Figure S2D**). This indicates that the TGFβ-1–treated MCF10A cells exhibit an Early- E/M state with ECM degradation decreased during G1 phase.

Together, these data indicate that the cell-cycle timing of invadopodia activity is not fixed, but changes with EMT progression.

### EMT progression shifts leader cell invasion from G2 to G1

During the 3D collective invasion of 4T1 spheroids, invadopodia are assembled solely at the front and sides of the leader cells, where they generate tracks that can subsequently be used by follower cells within the same invasive strand [16]. To determine whether the G2/G1 switch observed in 2D ECM degradation is reflected in leader-cell behavior, we embedded the control and TGFβ-1-treated 4T1-FUCCI (Early- and Late- E/M) spheroids in the dense (5 mg/ml) collagen I matrix, a condition that enforces MMP-dependent, invadopodia-mediated 3D invasion [41]. Compared to the spheroids containing Early-E/M cells, Late-E/M spheroids exhibited a more robust invasion (**Figure 2A, B**), characterized by an increase in the number of invasive strands (**Figure 2D**) and a two-fold increase of the invasion area (**Figure 2E**), indicating that Late-E/M have enhanced invasive capacity. Quantification of the leader cells in G1-phase (red) or G2 -phase cells (green) (**Figure 2C**) showed that the number of G1 leader cells has significantly increased with EMT progression from Early- to Late-E/M state. These findings suggest that the EMT-associated shift in invasion timing from G2 to G1 is reflected in leader-cell behavior and may be linked to increased invasive efficiency. This is consistent with previous studies showing that TGFβ-induced EMT promotes a more invasive phenotype in breast cancer cells [42,43].

**Figure 2.**
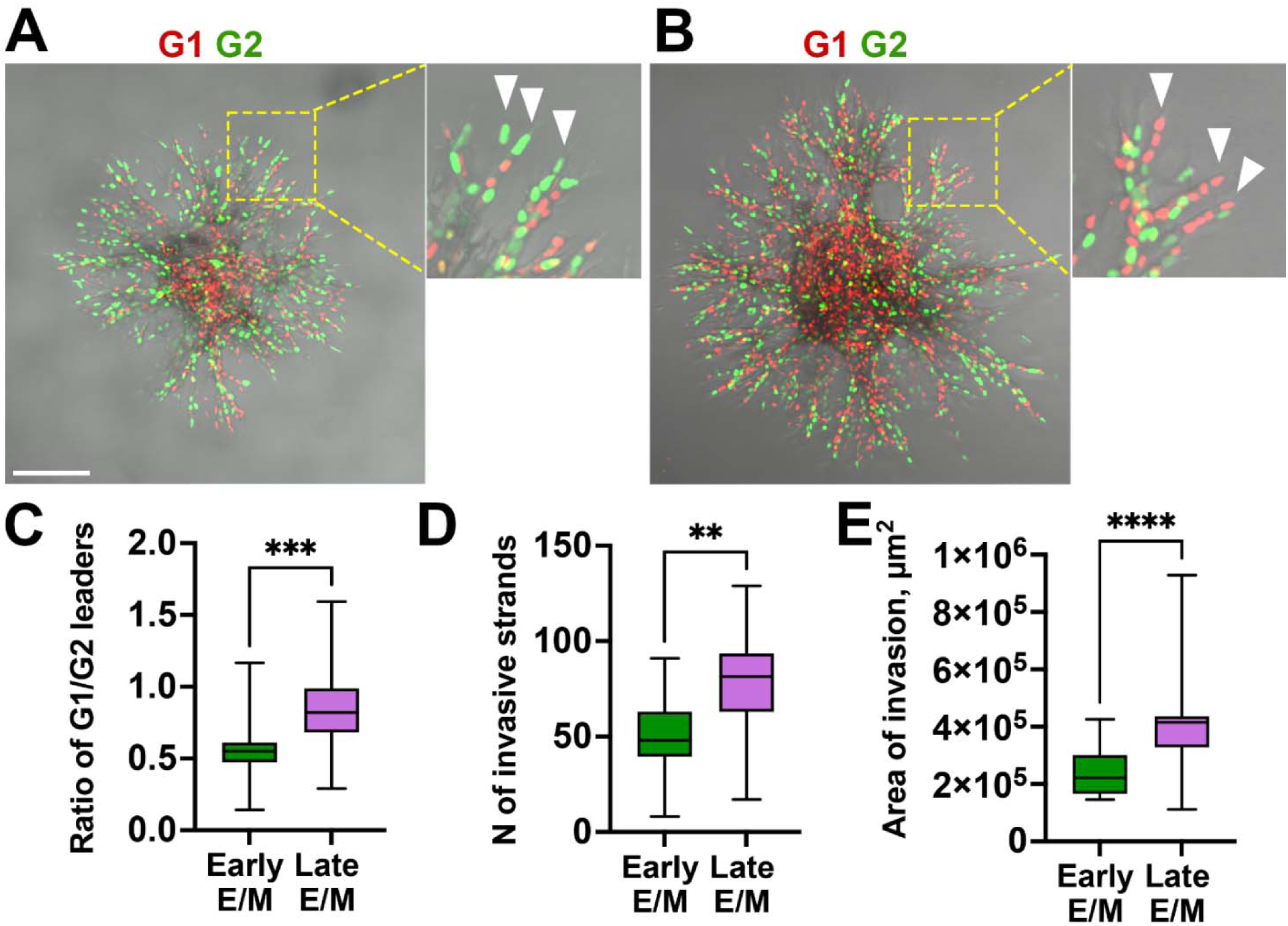
Most spheroid leaders are in G2 phase of the cell cycle in early E/M cells, and in G1 in late E/M cells. **A, B.** Spheroids of Early E/M (A) and Late E/M (B) cells, 48 hours post-embedding into high-density collagen. FUCCI G1 (red), FUCCI G2 (green). Scale bar: 200 µm. Dashed yellow boxes indicate regions magnified in inserts. White arrowheads mark leader cells at the tips of the invasive strands. **C.** G1/G2 leader cell ratio; ***P = 0.0005, Mann–Whitney U test. **D.** The number of invasive strands per spheroid in Early and Late E/M spheroids; **P=0.0078, Mann–Whitney U test. **E.** Total area of invasion; ****P <0.0001, Mann–Whitney U test. Data in (C–E) represent box-and-whisker plots showing median, interquartile range, and minimum/maximum values. N=3, 3-9 spheroids per N per condition.

Overall, these results indicate that the shift from Early to Late E/M state is associated with the shift of the invadopodia-rich leader cells from the G2 - to the G1-phase of the cell cycle.

### Bulk RNA-Seq reveals cell cycle-dependent expression of matrix remodeling genes and identifies FILIP1L as a master regulator of cell cycle-controlled invasion

To investigate the molecular mechanism responsible for the ECM degradation shifting from G2 phase in Early-E/M cells to G1 phase in Late-E/M cells, we established a 4-way sorting of the 4T1-FUCCI cells (**Figure 3A**). We classified cells as early- and late-E/M, and as G1 (red) or G2 (green). We then performed a transcriptomic profiling of all four classes using bulk RNA-Seq (**Figure 3A**). Pairwise differential gene expression analysis revealed significant transcriptional changes between Early- and Late- E/M across different cell cycle phases (**Figure 3B**). To further investigate the gene expression changes induced by TGFβ-1, we performed the pairwise comparisons of Early- versus Late- E/M states within the same cell cycle phase. Specifically, in the G2 phase, we identified 384 upregulated and 231 downregulated genes in Late- versus Early- E/M (**Figure 3B, top left**). Similarly, in the G1 phase, 470 genes were upregulated and 259 were downregulated (**Figure 3B, bottom right**). These results indicate that TGFβ-1-induced EMT shift had a substantial impact on the transcriptional landscape. Interestingly, TGFβ-1-induced EMT seems to be mediated primarily by the transcription factor Zeb1, while other classical EMT-associated transcription factors, including Twist1, Snai1, Snai3, and Zeb2, were not differentially expressed (**Figure 3B, top left and bottom right**).

**Figure 3.**
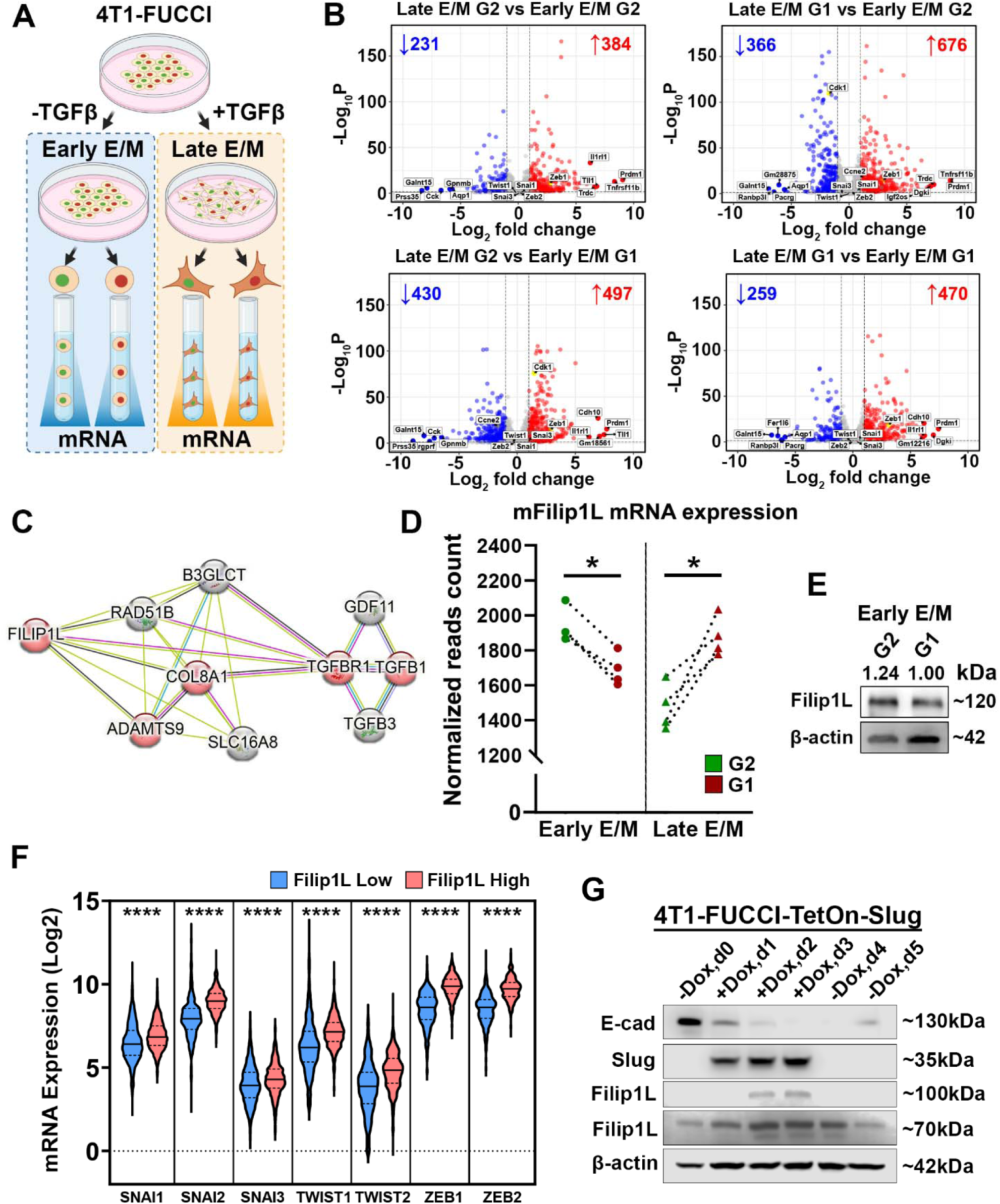
in Early E/M cells, FILIP1L expression is elevated in G2 phase, while in Late E/M cells, it is elevated in G1 phase of cell cycle. Increased FILIP1L correlates with worse prognosis in breast cancer patients. **A.** Schematic of multifactorial experimental workflow. 4T1-FUCCI cells were cultured with or without TGFβ, to achieve Late- or Early- E/M states, respectively. Cells were sorted into G1 or G2 phase bins based on FUCCI reporters, and mRNA was extracted for RNA-sequencing. **B.** Volcano plots showing differentially expressed genes between Early and Late E/M states across G2 and G1 phases of the cell cycle. Numbers in red and blue indicate significantly upregulated and downregulated genes, respectively (adjusted P < 0.05, log₂FC > 1 or < -1). **C.** STRING network of genes co-expressed with FILIP1L, highlighting interactions with EMT and TGFβ signaling-associated genes. **D.** FILIP1L mRNA expression in untreated (Early E/M) and TGFβ-treated (Late E/M), FACS-sorted 4T1-FUCCI cells. *P*□<□0.05 (*) Mann-Whitney U test. Dashed line represent the median. **E.** Western blot analysis of FILIP1L protein levels in untreated, early E/M, FACS-sorted 4T1-FUCCI cells. **F.** Violin plots of mRNA expression of EMT-TFS in TCGA samples, stratified by FILIP1L expression. FILIP1L-high group shows significantly higher levels of EMT-TFs (****P < 0.0001, Mann–Whitney U test). Dotted lines represent quartiles; the dashed line represents the median. **G.** Western blot showing upregulation of FILIP1L protein following Slug induction in 4T1-FUCCI-TetOn-Slug cells. Doxycycline was added at day 0, and lysates were collected at indicated timepoints. Patterns of expression of E-cadherin and Slug suggest EMT was completed and cells are in M state.

We next examined classes in which both EMT status and cell cycle phase contribute to gene expression changes. This analysis revealed 497 upregulated and 430 downregulated genes in the Late E/M G2 versus Early E/M G1 (**Figure 3B, bottom left**), and 676 upregulated and 366 downregulated genes in Late E/M G1 versus Early E/M G2 (**Figure 3B, top right**).

Using the Reactome database, we compiled a list of proteins and their corresponding genes associated with MMP activation (R-HSA-1592389), collagen degradation (R-HSA-1442490), and ECM degradation (R-HSA-1474228), alongside a curated set of invadopodia-related genes (**Figure S3, S4**). We next applied this gene set to our RNA-seq dataset to identify candidates exhibiting cell cycle–dependent expression.

In the Early- E/M cells, differential gene expression analysis revealed distinct shifts in ECM-associated transcriptional profiles between G1 and G2 phases (**Figure S3A**). Despite our prior evidence demonstrating that invadopodia-mediated ECM degradation of Early-E/M cells is cell cycle-dependent, the expression of core invadopodia components, including Cttn, Sh3pxd2a, Sh3pxd2b, Wasl, Arpc2, Mmp2, Mmp14, and Mmp9, remained unchanged across cell cycle phases (data not shown). Among matrix-degrading enzymes, only Mmp13 and Mmp10 were elevated in G1 (**Figure S3A**). Additional genes enriched in G1 included the matrix structural components Col1a2, Col6a1, and Col17a1 (**Figure S3A**). Notably, Nav3, encoding a microtubule-associated protein implicated in the negative regulation of ECM degradation and EMT [44,45], was upregulated in G2 (**Figure S3A**).

Upon transition to the Late-E/M state, a broader set of matrix remodeling genes began to exhibit cell cycle–dependent expression (**Figure S3B**). Mmp13 retained its upregulation in G1, whereas Mmp10 no longer showed significant phase-specific variation. Additional genes that became enriched in G1 included Mmp3 and several collagens associated with structural remodeling, such as Col5a3, Col7a1, Col12a1, Col1a1, and Col3a1. Adamts5 and Scube3, both of which are implicated in ECM degradation and modulation of TGF-β signaling, also showed increased expression in G1 (**Figure S3B)**. Among genes that acquired upregulation in G2 after cells transition in the Late E/M state are Adam8, which is responsible for fibronectin cleavage and modulating other proteases, and Tll3 (**Figure S3B)**. Thus, these results demonstrate that multiple ECM-degrading components show upregulation in either G1 or G2.

We also compared the expression of genes involved in MMP activation, collagen degradation, ECM remodeling, and invadopodia formation between cells in the same cell-cycle phase before and after TGFβ treatment (Early E/M vs. Late E/M) (**Figure S4**). This analysis revealed that an even greater number of matrix remodeling–related genes were differentially expressed between Early and Late E/M cells within the same cell-cycle phase.

Collagens (Col1a1, Col5a1, Col6a1, Col7a1, Col12a1, Col16a1, Col8a2) and laminins (Lamb3, Lamc2, Lama3) were enriched in Late E/M G1 cells (**Figure S4A**). Collagen I (Col1a1) and laminin β3 (Lamb3) are well-established EMT and metastasis markers [46,47], known to enhance adhesion to the basement membrane and facilitate migration through the ECM [46,48]. Matrix metalloproteinases (Mmp9, Mmp10, Mmp13, Mmp14) and ADAM/ADAMTS family proteases (Adam12, Adamts4, Adamts8, Adamts9) were also upregulated in Late E/M G1 cells compared to Early E/M G1 cells. These proteases promote ECM degradation and invadopodia-mediated invasion, consistent with the increased 3D invasiveness of Late E/M cells (**Figure 2**). In particular, MMP14 (MT1-MMP) plays a central role in ECM degradation at invadopodia. Fscn1 (Fascin), an actin-bundling protein essential for invadopodia protrusion, was also elevated in Late E/M cells. As expected, the epithelial marker Cdh1, encoding E-cadherin, was downregulated in Late E/M cells (**Figure S4A**).

Similar to Early vs Late cells in G1, in G2 cells MMPs (Mmp3, Mmp9, Mmp10, Mmp14) and ADAM family members (Adam12, Adam19, Adamts4, Adamts8) remain elevated in Late E/M G2 (**Figure S4B**). Laminins (Lamb3, Lama3, Lamc2) and collagens (Col1a1, Col4a5, Col5a1, Col16a1) are upregulated (**Figure S4B**), highlighting persistent ECM remodeling even in proliferative phases.

Together, this pattern suggests that Late E/M cells in both G1 and G2 phases are enriched for structural ECM and proteolytic machinery. This supports the data obtained in 3D collagen matrix when Late E/M cells were invading more actively (**Figure 2**), however, this does not allow to point out to the molecular machinery responsible for enriched number of G1 leaders in 3D (**Figure 2C**)

We hypothesized that if invadopodia components do not show cell cycle-dependent changes in expression, then a master regulator must exist that locally controls invadopodia function and is itself regulated by the cell cycle. To refine our analysis, we focused on genes associated with invasion in Early E/M G2 and Late E/M G1 cells, as these cell cycle phases were previously identified as the most invasive. Differential expression was assessed using a multi-group design implemented in DESeq2. This ensured that the differential expression analysis accounts for potential effects from both TGFβ treatment condition and cell cycle phase, isolating the effect of invasiveness on gene expression (**Table S1**). The result of this multivariate analysis showed two genes had significantly different gene expression (**Table S2**).

When corrected for the effects of cell cycle and TGFβ treatment on the RNA-seq dataset, Filamin interacting protein 1L, FILIP1L, was identified as the top candidate, as it had the most prominent difference of expression in the *invasive* group (G2 Early-E/M and G1 Late-E/M cells) compared to *non-invasive* group (**Table S2**). This understudied protein was never explored in relationship to invadopodia complex and we have proceeded to investigate it in detail.

### FILIP1L is co-expressed with EMT transcription factors in BRCA patients and linked to a mesenchymal phenotype

To explore the signaling context surrounding FILIP1L, we analyzed predicted protein–protein interaction network using the STRING database. This analysis revealed that FILIP1L is closely connected to several proteins involved in ECM organization and TGF-β signaling (**Figure 3C**). Notably, FILIP1L was linked to COL8A1, a basement membrane-associated collagen, which served as a central node connecting FILIP1L to components of the TGFβ-1 pathway. In addition, FILIP1L showed a link to ADAMTS9, one of the secreted metalloproteinases responsible for fibronectin degradation and collagen deposition [49,50].

Additionally, to determine whether Filip1L expression is cell-cycle dependent during EMT, we analyzed Filip1L expression in FACS-sorted 4T1-FUCCI cells untreated or treated with TGFβ (Early vs Late E/M). The analysis revealed that Filip1L mRNA levels were significantly higher in G2-phase cells (mAG sorted) compared to G1-phase cells (mKO2 sorted) in both Early E/M subpopulations (**Figure 3D**). However, upon TGFβ induction of Late E/M phenotype, Filip1L mRNA levels became higher in G1-phase cells (mKO2 sorted) (**Figure 3D**). Consistently, western blot analysis of Early E/M FACS-sorted 4T1-FUCCI cells confirmed elevated Filip1L protein expression in the G2 phase relative to G1 (**Figure 3E**). These results confirm that Filip1L is upregulated during the more “invasive” cell-cycle phase (G2 in Early E/M and GG1 in Late E/M) and are consistent with the invadopodia degradation assays of synchronized cells (**Figure 1B, I**).

The EMT is accompanied by the increased expression of proteolytic enzymes such as MMPs and ADAMs [51,52] and given that MMPs and ADAMs proteases are transcriptionally regulated by EMT-transcription factors (EMT-TFs) [53–55], we next investigated whether FILIP1L mRNA expression is correlated with that of EMT-TFs, ECM-remodeling proteases, and invadopodia-associated genes in patient-derived datasets. Analysis of the TCGA PanCancer Atlas breast cancer dataset, stratified into two cohorts based on the median FILIP1L expression level (FILIP1L-low vs. FILIP1L-high), revealed that FILIP1L expression positively correlates with the expression of key EMT-TFs, including members of the SNAI, TWIST, and ZEB family (**Figure 3F**). FILIP1L also showed positive correlation with several MMPs as well as members of the ADAM and ADAMTS families of proteases (**Figure S5A-B, Figure S5D-F**). FILIP1L expression was also positively associated with SH3PXD2A and SH3PXD2B, which encode the invadopodia scaffold proteins Tks5 and Tks4, respectively (**Figure S5C).**

To validate our previous data generated in TGFβ-1-induced EMT, we set out to investigate if FILIP1L expression is directly linked to EMT-transcription factor expression, which occurs downstream of TGFβ-1. We have established a 4T1 subline with inducible overexpression of Slug, 4T1-TetOn-Slug. Over the course of three days of doxycycline (+Dox) stimulation, as expected, Slug expression was increasingly upregulated, reaching the maximum value and plateauing on Day 2, maintain the value on Day 3 and immediately dropping on Day 4 when Dox- treatment was initiated (**Figure 3G**). Meanwhile, expression of the epithelial marker E-cadherin was increasingly downregulated Day 1-3, going back to its initial levels on Day 5. During the same period, FILIP1L protein levels were increasingly upregulated from Day 1 to Day 2, plateauing on Day 2, following the expression pattern of Slug (**Figure 3G**). Once Dox was removed, FILIP1L expression returned to its original level on Day 5 (**Figure 3G**). In summary, these findings support the idea that FILIP1L is integrated into the EMT regulatory network, is coordinated by both EMT state and cell-cycle phase, is associated with an invadopodia-rich phenotype, shows co-expression with EMT-associated transcription factors and ECM-remodeling proteases, and is transcriptionally regulated by Slug (**Figure 3G**).

### FILIP1L is a novel invadopodia component necessary for 3D spheroid invasion of E/M cells

Our analysis so far revealed a cell cycle phase-dependent role of FILIP1L in cancer cell invasiveness. We reasoned that FILIP1L may be a component in invadopodia and to test potential functional roles of FILIP1L, we examined its protein sequence. UniProt analysis revealed significant sequence homology between the N-terminal domain of FILIP1L and the N-terminus of cortactin-binding protein 2 (CortBP2; Pfam ID: PF09727) (**Figure 4A**). Additionally, amino acids 368–1128 of FILIP1L align with a region annotated as the Cortactin-Binding and Actin Dynamics Regulator domain (PANTHER database ID: PTHR23166; data not shown). Based on these findings, we hypothesized that FILIP1L is located in invadopodia.

**Figure 4.**
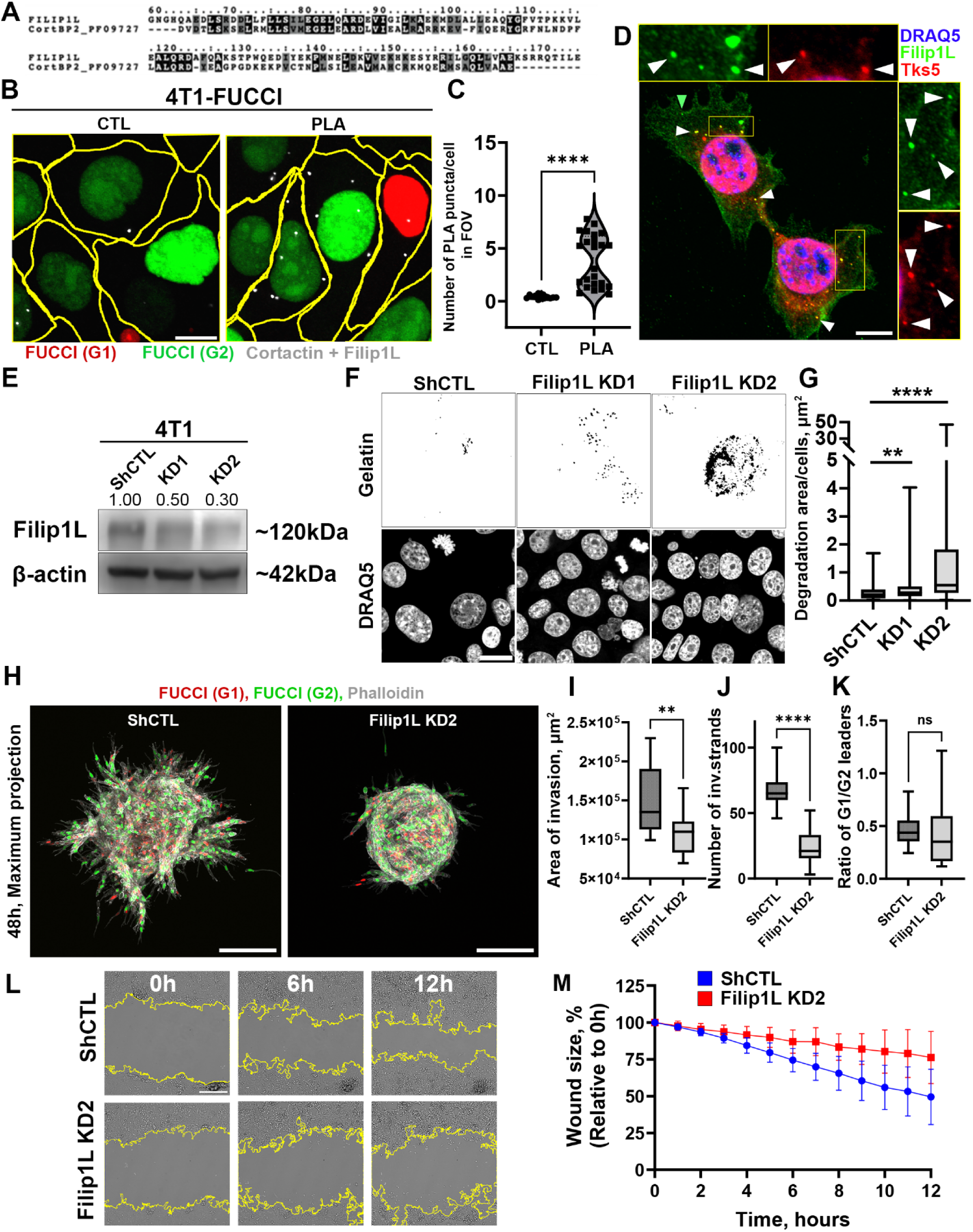
FILIP1L is a component of invadopodia. FILIP1L KD promotes ECM degradation, while inhibiting cell migration, reducing invasion of 3D spheroid. **A.** Protein sequence alignment of human FILIP1L (UniProt Q4L180, amino acids 59-174) and domain of cortactin-binding protein-2 (Pfam database accession PF09727). Identical residues are highlighted in black with white letters, conserved substitutions are shown in grey. **B.** Proximity ligation assay (PLA) detects presence of endogenous FILIP1L–cortactin interactions in 4T1-FUCCI cells. FUCCI G1 (Red), FUCCI G2 (Green), PLA puncta (white). Scale bar 10 μm. **C.** Number of PLA puncta per cell in negative control (no primary antibodies) and PLA (with primary antibodies).****P < 0.0001, Mann–Whitney U test. N=3, n=1, 3-11 FOVs per n. **D.** FILIP1L (green) and Tks5 (red) in 4T1 cells. White arrowheads show co-localizations of FILIP1L and Tks5, while green arrowheads show FILIP1L-only puncta. Nuclei were stained with DRAQ5 dye (blue). Inserts above and on the right of the image show magnified views of the boxed regions. Scale bar 20 μm. **E.** Western blot of 4T1 FUCCI- shCTL, FILIP1L KD1 and KD2 cells. FILIP1L levels were reduced by 50% in KD1 and 70% in KD2 cells. β-actin was used as the loading control. **F.** Binarized images of ECM degradation (black) in 4T1 FUCCI- shCTL, and FILIP1L KD1, KD2 cells. Nuclei labeled with DRAQ5. **G.** Quantification of ECM degradation per cell from images shown in F. (**P < 0.01; ****P < 0.0001; Mann–Whitney U test). Box-and-whisker plots showing median, interquartile range, and minimum/maximum values. N=3, n=3, 9-10 FOVs per n. **H.** Maximum intensity projections of 4T1 FUCCI- shCTL and FILIP1L KD2 spheroids embedded in 3D collagen I matrix for 48 h. FUCCI G1 (red) and G2 (green), phalloidin (white). Scale bars, 200 μm. **I-K.** Quantifications of (I) invasion area, (J) number of invasive strands, and (K) ratio of G1/ G2 leader cells. (ns P>0.05; **P < 0.01; ****P < 0.0001; Mann–Whitney U test). Box-and-whisker plots showing median, interquartile range, and minimum/maximum values. N=3, 5-6 spheroids per N per condition. **L.** Phase contrast images of the wound healing assay in 4T1-tdTomato-shCTL or FILIP1L KD2 cells. Scale bar 200□µm. **M.** Decrease in wound area size 0-12 hours post-scratch. Three independent biological replicates, 6 fields of view per replicate per condition. Data represented as the mean□±□SD. Statistical analysis was performed using two-way ANOVA with Geisser-Greenhouse correction (treatment × time interaction: F (12, 408) = 17.90, *P* < 0.0001; time: F (1.304, 44.35) = 114.5, *P* < 0.0001; treatment: F (1, 34) = 26.47, *P* < 0.0001). N=3, n=2, 6 FOVs per n per condition.

Using proximity ligation assay (PLA), we established that FILIP1L and cortactin are a part of the same complex in 4T1 cells (**Figure** □**4B, C**). Further, FILIP1L showed colocalization with the invadopodia-specific Tks5 protein (**Figure** □**4D**), supporting its localization to invadopodia in 4T1 E/M cells. Importantly, similar colocalization of FILIP1L and Tks5 was present in mesenchymal MDA-MB-231 cells (**Figure S6A**). FILIP1L also colocalizes with cortactin in mesenchymal MDA-MB-231 cells (**Figure S6B**). This makes a convincing point that FILIP1L is localized to invadopodia but leaves a question on whether FILIP1L is functionally required for invadopodia assembly or invadopodia function i.e. ECM degradation. To address this, we generated 4T1-FUCCI and 4T1-tdTomato sublines stably expressing control shRNAs (shCTL) or two independent FILIP1L shRNAs (KD1 and KD2), achieving approximately 50% and 70% knockdown efficiency, respectively (**Figure 4E**). Using these sublines, we performed invadopodia assay by culturing cells on fluorescently labeled gelatin over 20□hours. Quantification of gelatin degradation revealed that FILIP1L KD cells degraded significantly more ECM compared to control shCTL cells (**Figure 4F, G**), suggesting an increase in proteolytic activity in the absence of FILIP1L.

Interestingly, in 3D invasion assay, FILIP1L-KD2 spheroids displayed a reduced invasive capacity relative to FILIP1L-shCTL spheroids (**Figure 4H**), characterized by a smaller invasive area (**Figure 4I**) and fewer invasive strands (**Figure 4J**). Meanwhile, the ratio of G1/ G2-phase leader cells was consistent with Early E/M in 4T1 cells (**Figure 4K**). At a first glance, this result seemed to be inconsistent with increase in invadopodia-mediated ECM degradation. However, invasion consists of two states- invadopodia-rich ECM remodeling state, and migratory state during which cells translocate into the space generated by invadopodia. So, it is possible that FILIP1L-KD was favoring invadopodia state, while inhibiting the migratory state [23]. We tested this hypothesis using the wound healing assay, where FILIP1L-KD2 and FILIP1L-shCTL 4T1-tdTomato cells were cultured to 100% confluency, followed by a line-scratch and 12h of time-lapse imaging. As the wound is not filled with ECM, the wound closure was dependent solely on cell migration velocity. As suspected FILIP1L-KD cells exhibited significantly slower dynamics of wound closure compared to FILIP1L-shCTL, indicating lower velocity of cell migration (**Figure**□**4L, M**). In summary, while FILIP1L-KD shows a positive effect on invadopodia and ECM degradation levels, it had a contrasting, negative effect on cell migration in 2D and consequently, spheroid invasion in 3D.

Together, these findings identify FILIP1L as a critical invadopodia component required for productive invasion in 3D matrices, and directional cell migration, supporting its essential role in cancer invasion.

### FILIP1L KD impairs lung metastasis and lower FILIP1L expression correlates with better survival in patients

Efficient metastatic dissemination requires not only ECM degradation but also coordinated invasion that integrates ECM degradation with migration through the matrix to reach blood vessels. Given that FILIP1L knockdown increased ECM degradation (**Figure 4F, G**) but reduced migration and 3D spheroid invasion (**Figure 4H–J**), we sought to determine which of these effects better predicts metastatic outcome *in vivo*.

To investigate the role of FILIP1L in metastasis, 4T1-tdTomato sublines stably expressing control or FILIP1L shRNA (FILIP KD2) were inoculated into Balb/c mice. Cleared control tumors demonstrated colocalization of invadopodia marker Tks5 and FILIP1L in cancer cell next to a blood vessel (**Figure 5A**), suggesting that cells assembling invadopodia *in vivo* in order to metastasize express high levels of FILIP1L.

**Figure 5.**
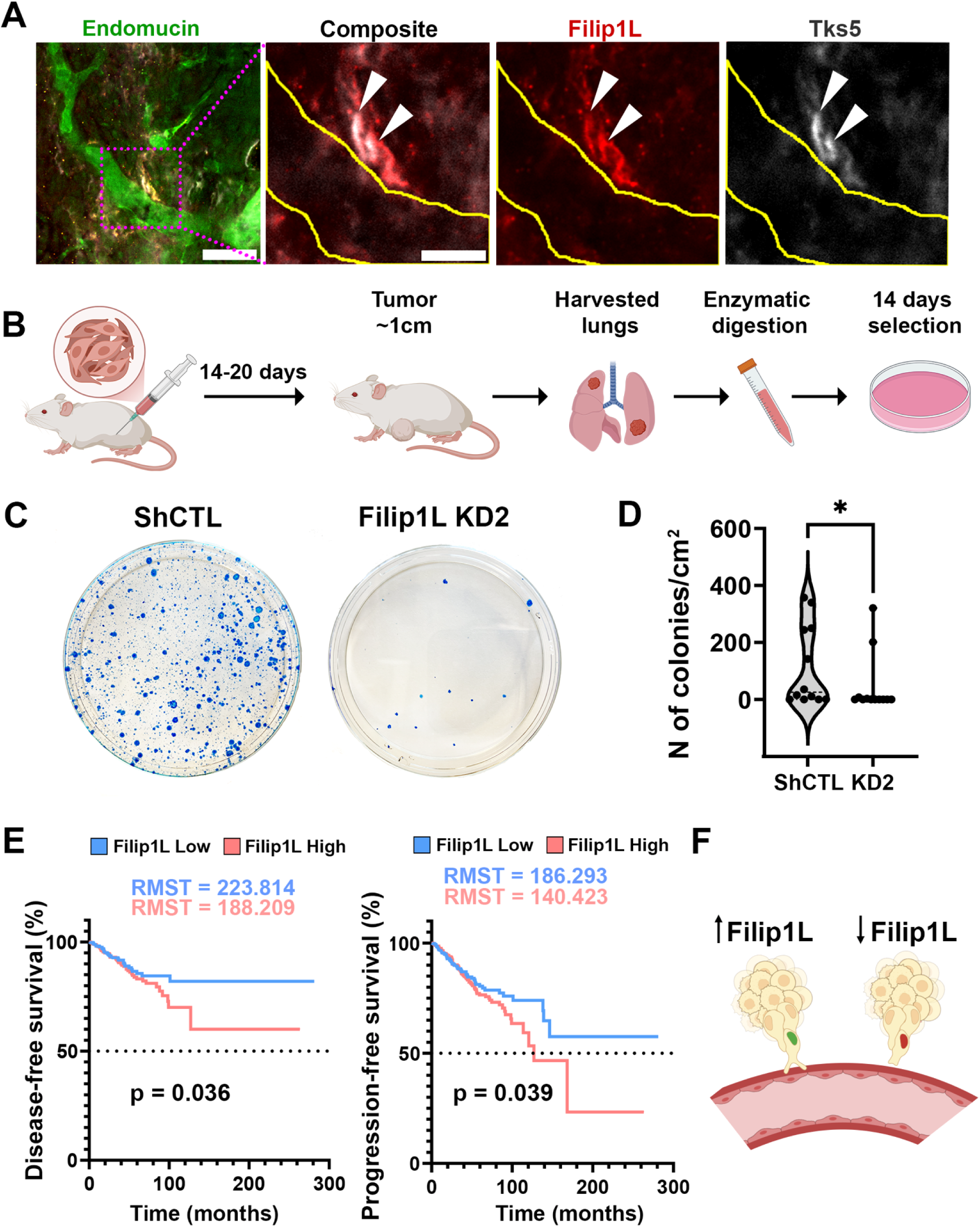
Lower FILIP1L expression correlates with lower metastatic colonization in mouse models and improved survival in breast cancer patients. **A.** Immunofluorescence image of cleared tumor labeled with blood vessel marker endomucin (green), FILIP1L (red), and Tks5 (white). The boxed region is shown at higher magnification in the right panels. Arrowheads indicate FILIP1L+, Tks5+ structures next to blood vessels. Yellow lines outline the blood vessel. Scale bars: 50 µm, 20 µm (boxed region). **B.** Schematic of the mouse clonogenic assay. **C.** Representative photographs of metastatic lung colonies from mice inoculated with control (shCTL) or FILIP1L knockdown (FILIP1L KD2) cells. **D.** Quantification of metastatic lung colonies per cm² for mice inoculated with shCTL and FILIP1L KD2 cells. Each dot represents one biological sample. P = 0.049. N=3, 2-5 mice per N. The dashed line is median. **E.** Kaplan–Meier curves of disease-free (left) and progression-free (right) in breast cancer patients stratified by FILIP1L expression levels (TCGA PanCancer Atlas). FILIP1L-low group correlates with a higher disease-free survival (*P* = 0.036), and a higher progression-free survival (*P* = 0.039). RMST, restricted mean survival time. **F.** Working model of invadopodia assembly and metastatic dissemination in early E/M cells: When leader cell is in G2, FILIP1L expression in increased and invadopodia degrades perivascular matrix. This allows leader cell and several followers to jointly enter the blood vessel, leading to metastasis. Created in BioRender.

Metastatic burden was assessed by clonogenic assay (**Figure 5B**). Despite the increased ECM degradation in FILIP1L KD cells *in vitro*, loss of FILIP1L markedly reduced the number of metastatic colonies in lungs (**Figure 5C, D**), with median values of 25.6 colonies/cm² in control tumors compared to 0.4 colonies/cm² in FILIP1L KD2 tumors **(Figure 5D**).

To understand the impact of FILIP1L expression on breast cancer patients, we have also assessed its prognostic relevance, classifying patients into FILIP1L-high and -low. Kaplan–Meier analysis based on median FILIP1L expression indicated that patients in the FILIP1L-low cohort exhibited a significantly longer restricted mean survival time (RMST).

The difference between RMST FILIP1L-low and high cohorts for the disease-free survival is ∼3.0 years (35.6 months), with the progression-free survival being ∼3.8 years (45.9 months) longer (**Figure 5E**). These findings suggest that FILIP1L is a crucial component for successful metastasis, serving as a functional link between EMT, cell cycle, invadopodia and tumor cell dissemination in human patients.

Together, these findings indicate that FILIP1L is required not simply for matrix proteolysis, but for the coordination between invadopodia and migration states, which constitutes productive invasive behavior [23,24]. In this context, increased ECM degradation is not sufficient for metastasis, suggesting that dissemination of hybrid E/M cells depends on proper coordination between degradative and migratory states. These data therefore position FILIP1L as a key regulator of 3D invasion and metastatic dissemination in hybrid E/M breast cancer cells. When FILIP1L is present and upregulated during G2, invasion and metastasis occur; in contrast, low levels of FILIP1L impair invasion, consequently preventing metastasis (**Figure 5F**).

## Discussion

In this study, we identify EMT state as a determinant of when breast cancer cells invade during the cell cycle. We show that in Early E/M cells invadopodia-mediated ECM degradation and invasion occur predominantly in G2 phase of the cell cycle, meanwhile progression towards Late E/M and fully M states shift invasion to G1 phase. This pattern was observed not only in 2D degradation assays, but also in 3D spheroid invasion, where Early E/M leader cells were enriched in G2, while Late E/M leader cells were preferentially found in G1. Thus, EMT progression does not simply alter the magnitude of invasion, but reorganizes when invasion occurs during cell cycle. Importantly, this data is in alignment with our previously published report of fully M human breast cancer cell lines MDA-MB-231 and BT-549 degrading ECM, invading in 3D and intravasating *in vivo* during G1 phase [25].

Important finding of this study is the identification of a G2-associated invasive state in Early E/M cells. Because hybrid E/M state is increasingly recognized as highly plastic and metastatic, capable of cooperative metastasis, G2-associated invasion may represent a specialized strategy that allows these cells to retain epithelial features while simultaneously engaging matrix remodeling and collective behavior. In this way, G2 invasion may be a defining trait of hybrid-state biology. More broadly, these results suggest that proliferative status and the ability to invade ECM are linked in hybrid E/M state.

Interestingly, while the TGFβ-treatment is a well-recognized strategy for achieving EMT [56], it resulted in Early E/M cells transitioning to a later E/M state, but not to M, as the E-cadherin expression was preserved. As previously reported, and confirmed by our RNA sequencing data, TGFβ-induced EMT is regulated by the transcription factor Zeb1 [35]. However, overexpression of a different EMT-related transcription factor, Slug, led to a complete EMT transition to M state, judged by the loss of E-cadherin expression. As previously hypothesized [10], we here demonstrate that different EMT programs, driven by specific transcription factors, can result in distinct and stable states along the EMT spectrum. However, independent of the specific transcription factor involved, we show that the shift invadopodia function predominantly occurring in G2 in Early E/M vs in G1 phase in Late E/M, or M cells, appear to be a characteristic of progression along the EMT axis. Our data is further strengthened by the results of the TGFβ-treatment in normal epithelial (E) cell line MCF10A. While untreated cells have shown almost no ECM degradation, treated cells transition to an Early E/M state, with an increase in vimentin and maintained expression of E-cadherin and with G2-dominant invadopodia-mediated ECM degradation.

Our findings contribute to the growing recognition that EMT is not a linear or uniform process, but rather a multidimensional spectrum governed by distinct transcriptional programs. While EMT can be driven by various master regulators, such as Slug or Zeb1, they seem to activate overlapping but non-identical gene expression networks. For instance, Slug-driven EMT and TGFβ-induced EMT (mediated by Zeb1) represent divergent regulatory trajectories: the former leads to a complete E-cadherin downregulation, degradative activity predominantly occurring in G1, and in an overall increase in ECM degradation, whereas the latter results in only partial E-cadherin downregulation, degradation occurring in G1, but without an overall increase in degradation levels. This suggests that EMT progression may not follow a single axis of EMT differentiation but instead spans a multidimensional landscape, where cells can occupy hybrid or intermediate states depending on the combination and levels of transcription factors expressed. In fully M cells such as MDA-MB-231, multiple EMT-TFs are co-expressed at high levels, potentially representing a terminal or stabilized M state [25].

Using RNA-seq of cell-cycle-sorted Early and Late E/M cells, we identified FILIP1L as an EMT- and cell-cycle-regulated candidate whose expression matches the invasive phase of each state: enriched in G2 in Early E/M cells and in G1 in Late E/M cellscells.FILIP1L colocalized with Tks5 and cortactin, consistent with its role as an invadopodia component. Aligning with our findings on the upregulation of FILIP1L in the more invasive Late E/M cells [57], the authors observed that purely epithelial (E) human breast cancer MCF7 cells did not express FILIP1L, whereas purely M human breast cancer MDA-MB-231 cells had a high FILIP1L expression [57]. Interestingly, same report suggested that FILIP1L may localize to mitochondria, something we did not detect, and could be due to the cells being plated in the absence of ECM layer.

A previous study in an ovarian cancer model identified FILIP1L as a Wnt/β-catenin signaling inhibitor associated with EMT [58]. Doxycycline-inducible FILIP1L expression was shown to reduce MMP levels and suppress invasion. Additionally, lower FILIP1L expression correlated with poorer disease-free and overall survival in ovarian cancer patients [59]. Finally, FILIP1L expression was inversely correlated with Slug expression in ovarian cancer patients [59]. However, the role of FILIP1L in breast cancer or relationship to the cell cycle-control of invasion had not previously been explored.

Our localization studies further support a role for FILIP1L in invadopodia. FILIP1L associated with cortactin by proximity ligation assay and colocalized with Tks5 in 4T1 cells, with similar localization observed in mesenchymal MDA-MB-231 cells. This data strongly supports the conclusion that FILIP1L is an invadopodia-associated protein. However, our functional findings indicate that FILIP1L is not a simple positive regulator of matrix degradation. In contrast, FILIP1L loss increased gelatin degradation while simultaneously reducing migration, impairing 3D spheroid invasion, and markedly decreasing metastatic colonization in vivo. This combination of phenotypes suggests that FILIP1L supports productive invasion by coordinating the balance between degradative and migratory states required for efficient movement through 3D matrix.

We further demonstrate that FILIP1L is upregulated upon Slug overexpression *in vitro* and co-expressed with key EMT transcription factors such as Slug, Snail, Twist, and Zeb in patient data. This indicates that FILIP1L is transcriptionally regulated by Slug and is part of a mesenchymal gene expression program in breast cancer. Interestingly, in contrast to ovarian cancer, breast cancer patients with low FILIP1L expression in TCGA datasets had better disease-free and progression-free survival.

The discrepancy between FILIP1L’s role in breast and ovarian cancer may stem from differences in their corresponding metastatic routes. Specifically, while breast cancer cells require active ECM degradation and primarily metastasize through hematogenous and lymphatic routes [60], ovarian cancer cells may passively disseminate into the peritoneal cavity due to the absence of anatomical barriers [61].

In summary, our findings reveal that the Early E/M states possess a unique capacity to assemble invadopodia and invade during the G2 phase of the cell cycle thus demonstrating that proliferation and invasion programs are coupled in hybrid E/M cells. Moreover, We further identify FILIP1L as an invadopodia-associated, EMT-and cell-cycle-regulated factor that supports productive invasion and metastatic colonization. Importantly, our results highlight the tissue-specific role of FILIP1L, suggesting that in breast cancer it functions as a metastasis promoter and is a marker of the mesenchymal phenotype.

## Acknowledgements

We thank the following funding sources for their support: NIH NCI R01 CA230777; American Cancer Society Research Scholar Grant 134415-RSG-20-34-01-CSM, DOD BCRP Breakthrough Award BC230197, PA-CURE, Temple STEM Research Award and WW Smith Charitable Trust awards to B.G. We thank Dr. Wenjun Guo from Albert Einstein College of Medicine, Bronx, NY for helpful discussions. We also thank Amir Yarmahmoodi and the Flow Cytometry Core Facility at the Lewis Katz School of Medicine at Temple University for assistance with cell sorting.

## Author contributions

Design of the study: BG, EB; Data Acquisition: EB, AJ. Data Analysis and interpretation: EB, AJ, BG, GA and ALK. Supervision: BG; Fund Acquisition: BG; Manuscript writing: EB, BG.

## Data Availability

All mRNA sequencing data have been uploaded to the GEO database.

## Materials and methods

### Ethics statement

Experiments involving mice (*Mus musculus*) were conducted according to the NIH regulations and approved by the Temple University (IACUC protocol #5072)

### Cell lines and culturing conditions

Early E/M murine mammary carcinoma cell line 4T1 was a gift from Dr. Fred R. Miller at the Karmanos Cancer Center. Cells were maintained in high-glucose Dulbecco’s Modified Eagle Medium (11965092, Gibco, Grand Island, NY, USA) (4.5 g/L D-glucose, L-glutamine), supplemented with 10% fetal bovine serum (S11150, Atlanta Biologicals, Norcross, GA, USA) and 50 U/mL penicillin – 50 µg/mL streptomycin (Gibco, Grand Island, NY, USA). Cultures were incubated at 37□°C with 5% CO₂ for up to 30 days.

A stable 4T1-FUCCI subline was generated through the sequential lentiviral transduction with red mKO2-hCdt1 (amino acids 30/120) and green mAG-hGem (amino acids 1/110) FUCCI constructs. The mKO2-hCdt1 and mAG-hGem constructs were obtained from the RIKEN BioResource Center, Gene Engineering Division, Japan. HEK293T cells (∼60% confluency) (a gift from the George Smith laboratory, Temple University) were transfected using Lipofectamine 3000: 36□µl reagent was mixed with 800□µl OptiMEM; 24□µg total DNA (6□µg pCMV-VSV-G-RSV-REV, 6□µg pCAG-HIVgp, 12□µg FUCCI) and 48□µl P3000 were mixed with 800□µl OptiMEM. Solutions were incubated separately for 5□min, then combined and incubated for 20□min at RT before adding ∼1□ml per 10□cm dish in antibiotic-free DMEM +10% FBS. Virus was collected 48h post transfection and treated with Lenti X concentrator according to the manufacturer’s instructions (631231, Takara Bio, Göteborg, Sweden). Adherent 4T1 cells 30-40% confluency were infected with 24 µl 1:10 dilution of virus into serum-free DMEM in the presence of 10 µg/ml polybrene in 10cm dish. To ensure the presence of both mKO2 and mAG in the cells, top 10% bright mKO2⁺ cells were first selected by FACS, allowed to recover and expand for 7 days, and subsequently sorted again to select for top 10% bright mAG⁺ population.

Stable expression of the reverse tetracycline transactivator (rtTA) was achieved by co-transfecting 4.0 × 10□ HEK293T cells with 3000 ng total DNA per well comprising rtTA-N144, pMD2.G, and pCMVR8.74 at a ratio of 1.8:1.0:2.2 using Lipofectamine™ 3000 Transfection Reagent (L3000008, Thermo Fisher Scientific, Carlsbad, CA, USA), according to the manufacturer’s instructions. rtTA-N144 was a gift from Andrew Yoo (Addgene plasmid # 66810; http://n2t.net/addgene:66810 RRID:Addgene_66810) [62]. pMD2.G and pCMVR8.74 were gifts from Didier Trono (Addgene plasmid #12259; http://n2t.net/addgene:12259; RRID:Addgene_12259 (unpublished) and Addgene plasmid #22036; http://n2t.net/addgene:22036; RRID:Addgene_22036 (unpublished)). Lentiviral supernatant was collected 48 h post-transfection, filtered, and used to transduce 2.0 × 10□ 4T1 or 4T1-FUCCI cells in the presence of 10 µg/ml polybrene (TR-1003-G, Sigma-Aldrich, St. Louis, MO, USA). Cells were allowed to recover for 48 h prior to selection with 400 µg/ml hygromycin B (10687010, Thermo Fisher Scientific, Carlsbad, CA, USA) for 7 days.

To generate Slug-expressing cells, HEK 293T cells were co-transfected with pTK-Slug, pMD2.G, and pCMVR8.74 at the same ratio and total DNA concentration as described above. pTK-Slug was a gift from Bob Weinberg (Addgene plasmid #36986; http://n2t.net/addgene:36986; RRID:Addgene_36986) [63]. Viral supernatant was collected and used to transduce 4T1-rtTA and 4T1-FUCCI-rtTA cells. Following recovery, transduced cells were subjected to single-cell cloning by limiting dilution in 96-well plates under continuous selection with hygromycin B. After 7 days, wells containing individual colonies were supplemented with medium containing 2 µM doxycycline (D5207-1G, Sigma-Aldrich, St. Louis, MO, USA) to induce Slug expression. Colonies exhibiting a transition from epithelial to mesenchymal morphology were identified, expanded in doxycycline-free medium, and used for subsequent experiments.

FILIP1L knockdown cell lines (-KD1 and -KD2) and corresponding control (shCTL) lines were generated by transducing 4T1, 4T1-FUCCI and 4T1-tdTomato cells with lentiviral particles at a multiplicity of infection (MOI) of 3. Lentiviral vectors contained shRNA targeting FILIP1L (Clone ID: TRCN0000346916, Target Sequence: TGATAACTACTGAGGATAATA; Clone ID: TRCN0000346987, Target sequence: GAGTATCTGGAACTAACTATG) and non-target shRNA (shCTL) in the pLKO.5-puro vector backbone (MISSION library, Sigma-Aldrich). Following transduction, cells were selected with 4 µg/ml puromycin (ICN10055210, Irvine, CA, USA) for 7 days. Knockdown efficiency was validated by western blot analysis.

### Invadopodia degradation assay

Gelatin was labeled with Alexa-405-NHS ester for fluorescence, and 35□mm glass-bottom dishes or 24-well glass-bottom plates (MatTek Corporation, Ashland, MA, USA) were coated with Alexa-405-gelatin following a previously established protocol [64]. Briefly, MatTek dishes were pre-treated with 1N HCl (7647-01-0, Fluka, Geneva, Switzerland) for 10 minutes, followed by coating with 50□µg/mL Poly-L-Lysine (P8920, Sigma-Aldrich, St. Louis, MO, USA) for 20 minutes and multiple PBS washes. A 0.2% fluorescently labeled gelatin solution was prepared by diluting 2% gelatin and mixing with thawed dye-labeled gelatin aliquots for 10 minutes. After removal of gelatin, dishes were crosslinked with 0.2% glutaraldehyde (G5882, Millipore Sigma, Burlington, MA, USA) on ice for 15 minutes, then quenched with 5□mg/mL sodium borohydride (452882, Sigma-Aldrich, St. Louis, MO, USA) for 15 minutes. Following additional PBS washes, dishes were disinfected with 70% ethanol and finally incubated with Penicillin/Streptomycin in PBS. All steps were performed under low light conditions, and prepared dishes were stored at 4°C in the dark.

Cells were seeded at densities of 400 000 cells (35 mm glass-bottom dish) or 60 000 cells/well (24-well glass-bottom plate) and allowed to degrade the gelatin coating for 20h. Cells were then washed twice with PBS (Gibco, Grand Island, NY, USA) at RT and fixed (Fixation procedure is described in the Immunofluorescence section).

Images of gelatin degradation and cell nuclei were collected using a laser scanning confocal microscope (FV1200, Olympus, Tokyo, Japan) equipped with a 60X objective lens (UPLSAPO60XS, 1.35 NA, Olympus, Tokyo, Japan). Z-stack images were captured with 0.25 µm step sizes. Gelatin channel images were then analyzed using a custom macro in Fiji, first compiling the Z-stack slices into a Max Intensity projection, and then thresholding the signal. Degradation puncta were identified using the Particle Analysis tool. To account for differences in cell density across the fields of view, the total degradation area was normalized by dividing it by the number of cells in the same field, with the number of cells determined based on the nuclei staining (DAPI, 62248 or Draq5, 62251, Thermo Scientific, Rockford, IL, USA).

### Cell cycle synchronization

In the 6 well plate, 300 000 cells were seeded 24h prior to the synchronization. To synchronize, cells were incubated for 24 hours with 10 µM lovastatin (G1 phase synchronization) (PHR1285, Sigma-Aldrich, St. Louis, MO, USA), or 1 µg/mL mitomycin C (G2 phase synchronization) (11435, Cayman Chemicals, Ann Arbor, MI, USA). Control cells were incubated with DMSO (Cayman Chemicals, Ann Arbor, MI, USA) for 24h.

### Immunofluorescence

Immunofluorescent labeling of cells in 2D was performed after fixation with 4% paraformaldehyde (PFA, Alfa Aesar, Ward Hill, MA, USA) for 10 minutes RT. After fixation, cells were permeabilized with 0.1% Triton X-100 (Calbiochem, Merck KGaA, Darmstadt, Germany) at RT for 5 minutes, blocked with 1% FBS/5% BSA (Sigma-Aldrich, St. Louis, MO, USA) in PBS (Gibco, Grand Island, NY, USA) for 3h RT. After blocking, cells were incubated with primary, overnight 4°C in blocking, and secondary antibodies, 1h RT in blocking. All materials are described in **Table S3**.

### Western blots

Cells were lysed in ice-cold RIPA buffer (Teknova, Hollister, CA, USA) supplemented with protease inhibitors (complete cocktail, Roche) and phosphatase inhibitors (Halt cocktail, 78420, Sigma-Aldrich, St. Louis, MO, USA). Protein samples (20-30 µg per lane) were separated by SDS-PAGE and transferred onto a polyvinylidene difluoride membrane (Immobilon, Merck KGaA, Darmstadt, Germany). Membranes were blocked with 5% BSA in TBST for 3 hours at room temperature before overnight incubation at 4°C with primary antibodies. The following day, membranes were washed 3 times with TBST (10 min wash) and incubated for 1 hour at room temperature with HRP-conjugated anti-mouse or anti-rabbit IgG, 1:5000 dilution (Cell Signaling Technologies, Danvers, MA, USA) diluted in 5% non-fat milk/TBST. Protein bands were detected using chemiluminescence reagents and imaged with a blot scanner (C-DiGit, LI-COR Biosciences, Lincoln, NE, USA). Two chemiluminescent substrates were used: WesternBright ECL, (K-12045-D20, Advansta, Menlo Park, CA, USA) and Supersignal West Femto (34094, Thermo Scientific, Rockford, IL, USA). All materials are described in **Table S4**.

### Fluorescence-activated cell sorting (FACS) and bulk RNA sequencing

Early E/M (4T1-FUCCI) and Late E/M (4T1-FUCCI treated with TGFβ-1) cells were detached using 0.05% trypsin-EDTA, neutralized with complete medium, and resuspended in Accumax Cell Aggregate Dissociation Medium (00-4666-56, Thermo Fisher Scientific, Waltham, MA, USA) according to the manufecturer’s instructions and placed on ice.

Prior to sorting, cell suspensions were filtered through a 70□µm mesh strainer to eliminate aggregates. Fluorescence was detected using a BD FACSAria IIµ. Sorted cells were collected directly into tubes containing ice-cold, RNase- and DNase-free 1× PBS. Samples were centrifuged at 1200 rpm for 5□min at 4□°C, and the cell pellets were immediately flash-frozen and stored at –80□°C for downstream RNA extraction.

Frozen cell samples were shipped to Genewiz (Azenta Life Sciences, Burlington, MA, USA) for RNA extraction, library preparation, and next-generation sequencing using their RNAseq Transcriptome Profiling service. Quality control, library construction, and sequencing were performed according to the standard protocols provided by Genewiz. Downstream analysis was conducted in R using the DESeq2 package.

Further analysis was performed to go beyond simple two-group comparisons by incorporating multiple independent variables into the design formula, allowing identification of genes associated with the invasive phenotype: design = ∼ Invasiveness + Treatment + Color

### Proximity ligation assay

Proximity ligation assay (PLA) was performed using Duolink In Situ PLA Probe Anti-Mouse MINUS (DUO92004-30RXN, Sigma-Aldrich, St. Louis, MO, USA) and Duolink In Situ PLA Probe Anti-Rabbit PLUS (DUO92002-30RXN, Sigma-Aldrich, St. Louis, MO, USA) according to a manufacturer’s instructions. Briefly, 4T1 or 4T1-FUCCI cells were cultured on the gelatin-coated Mattek dish. At 24h, cells were fixed using 4% PFA for 10 min RT and permeabilized with 0.1% Triton-X for 5 min RT, blocked with blocking reagent provided in Duolink In Situ PLA Probe for 30min at 37C° and incubated with primary antibodies: mouse anti-Cortactin Clone 4F11 (05-180-I-100ul, Millipore Sigma, Burlington, MA, USA) and rabbit anti-FILIP1L (ab151331, Abcam, Cambridge, UK), diluted to 2.5 µg/ml in blocking reagent and incubated for 1h at 37C°. Cells were washed with Duolink In Situ Wash Buffer A, Fluorescence (DUO82049, Sigma-Aldrich, St. Louis, MO, USA) 2 x 5min RT and incubated with Duolink In Situ PLA Probe Anti-Mouse MINUS and Duolink In Situ PLA Probe Anti-Rabbit PLUS diluted 1:5 in diluent provided in the kit, 1h at 37C°. Cells were washed with Buffer A, 2 x 5min RT. Probes were ligated 30min at 37C° and amplified 1hr 40min at 37C° (Duolink In Situ Detection Reagents FarRed (DUO92013-30RXN, Sigma-Aldrich, St. Louis, MO, USA). PLA puncta were visualized using confocal microscope with oil 60x objective and the number of PLA puncta was normalized to the number of cells in the field of view (FOV).

### 3D Spheroid invasion assay

3D spheroids were generated using a suspension culture protocol adapted from [16]. Cells were trypsinized and resuspended at a density of 2000 cells/mL in spheroid formation medium composed of complete DMEM/F12 (5% horse serum (16050-122, Gibco, Grand Island, NY, USA), 0.5 µg/ml hydrocortisone (H0888, Sigma-Aldrich, St. Louis, MO, USA), 20 ng/ml hEGF (PHG0311L, Thermo Fisher Scientific, Waltham, MA, USA), 10 µg/ml insulin (91077C, Sigma-Aldrich, St. Louis, MO, USA), 100 ng/ml cholera toxin (C8052, Sigma-Aldrich, St. Louis, MO, USA) 1% penicillin/streptomycin (15140122, Thermo Fisher Scientific, Waltham, MA, USA) supplemented with 0.25% methylcellulose (M6385, Sigma-Aldrich, St. Louis, MO, USA). A volume of 200□µl (400 cells) was seeded per well into non-adherent round-bottom 96-well plates (Costar, cat. no. 3788). Plates were sealed with tape and centrifuged at 1000□rpm for 5□min at room temperature to allow cells to settle into a uniform sheet at the bottom of each well.

Following centrifugation, plates were transferred to an orbital shaker placed inside a humidified incubator set to 37□°C and 5% CO₂. The cells were agitated at ∼60□rpm for 2□h to facilitate initial compaction. The spheroid formation medium was then removed using a multichannel pipette, and 200□µl of cold spheroid growth medium, complete DMEM/F12 supplemented with 0.25% methylcellulose and 1% Matrigel (Corning, NY, USA, Cat #356234), was added to each well. Plates were returned to the incubator and maintained under static conditions for 48□h.

At 48□h, spheroids were collected and embedded in 30□µl of 5□mg/mL rat tail Collagen I (Corning, NY, USA, Cat # 354249) prepared according to the alternate gelation protocol. The collagen mixture containing individual spheroids was placed into custom-fabricated spheroid imaging devices (SIDs) [27]. Collagen was polymerized at 37□°C for 30□min, after which pre-warmed culture medium was added to each well to fully cover the gel.

### Wound healing assay

Six-well plates were pre-coated with 50 µg/ml poly-L-lysine (P8920, Sigma-Aldrich, St. Louis, MO, USA) for 20 min, followed by air drying. Cells were seeded and cultured until a confluent monolayer formed. A cross-shaped wound was then introduced using a 10 µl pipette tip. Time-lapse imaging was conducted using Nikon Eclipse Ti2 microscope using a 10x objective (Nikon Corporation, Tokyo, Japan), covering 1.33x1.33mm FOV. Images were acquired at 1 h intervals over a 12 h period. Wound size in the FOV was manually analyzed with Wand (tracing) tool in Fiji. Wound sizes were then normalized to the size of wound at 0 h.

### Clonogenic lung metastasis assay

To generate orthotopic mammary tumors, 200 000 cells were suspended in 100 μL of 20% collagen I in PBS and injected into the mammary fat pad of 7-week-old female BALB/c mice. Mice were euthanized 14–20 days after injection, once primary tumors reached approximately 1 cm in diameter, and the tumors and lungs were harvested. Lungs were minced, digested in a collagenase type IV/elastase cocktail (Worthington Biochemical, Lakewood, NJ, USA, LS004186, LS002274), and passed through a 70μm cell strainer. The resulting cell suspension was centrifuged, resuspended, and plated in DMEM supplemented with 10% FBS, 1% penicillin/streptomycin and 0.25 µg/ml Amphotericin B. To select for 4T1 cells, the medium was supplemented with 60 µM 6-thioguanine (15774, Cayman Chemicals, Ann Arbor, MI, USA). After 14 days of culture at 37 °C in 5% CO2, colonies were fixed with cold methanol, stained with 0.03% (w/v) methylene blue (457250, Sigma-Aldrich, St. Louis, MO, USA), and counted.

### CUBIC clearing of tumors

Tumors harvested from BALB/c mice were fixed overnight in 4% paraformaldehyde (PFA), cut into approximately 1 × 0.5 × 0.5 cm sections, and cleared using the CUBIC method. Tumor sections were first incubated in 50% CUBIC reagent 1 overnight at 37 °C, followed by 100% reagent 1 for 2–3 days at 37 °C. Reagent 1 consisted of urea (25% [w/w]), N,N,N′,N′-tetrakis(2-hydroxypropyl)ethylenediamine (25% [w/w]), and Triton X-100 (15% [w/w]) in distilled water [65]. Samples were then washed in PBS overnight and blocked in 5% BSA in PBS overnight at RT.

Tumor sections were immunolabeled with primary antibodies against endomucin (1:100; Santa Cruz Biotechnology, sc-65495), Tks5 (1:75; EMD Millipore, MABT336), and FILIP1L (1:100; Thermo Scientific, PA5-60251) diluted in blocking buffer for 48 h at 4 °C. After primary antibody incubation, samples were washed 3 times in PBS for 2 h each and then incubated with secondary antibodies diluted 1:100 in blocking buffer overnight at 4 °C. Samples were subsequently washed 3 times in PBS for 2h each and subjected to refractive index matching by incubation in 50% CUBIC-R+(M) tissue-clearing reagent (TCI, T3741) overnight at 37 °C, followed by 100% CUBIC-R+(M) for 48h at 37 °C. Cleared tumor sections were imaged within 7 days.

## Abbreviations

2D: Two-dimensional
3D: Three-dimensional
ECM: Extracellular matrix
EMT: Epithelial-mesenchymal transition
E/M: Epithelial/mesenchymal (hybrid state)
FBS: Fetal bovine serum
FILIP1L: Filamin A Interacting Protein 1-like
MET: Mesenchymal-to-epithelial transition
PBS: Phosphate-buffered saline
RNA-seq: RNA sequencing
RT: Room temperature
TGFβ: Transforming growth factor beta

## Supplementary Materials

**Supplementary Table 1. DESeq2 design setup**

**Supplementary Table 2. Result of Multivariate DESeq2 analysis**

**Supplementary Table 3. Materials for immunofluorescence**

**Supplementary Table 4. Materials for Western blots**

**Suplementary Figure 1. Induced overexpression of Slug leads to increases in invadopodia-mediated ECM degradation and in the number of G1 cells.**

**A.** Binarized images of invadopodia degradation (black puncta) in 4T1-FUCCI-TetOn-Slug cells cultured on fluorescent gelatin for 16□h, synchronized with DMSO, Lov, or MitC. Prior to plating on gelatin, cells were exposed to -Dox or +Dox for 3 days. Nuclei were stained with DRAQ5 (magenta). Scale bar, 20□µm.

**B.** Fluorescence overlay images of 4T1-FUCCI-TetOn-Slug cells in G1 (red) and G2 (green), imaged on day 3 after addition of doxycycline. Scale bar, 100□µm.

**C.** Quantification of cell cycle distribution in -Dox d3 and +Dox d3 cells from panel B. Data represent mean ± SD. N=3, n=3, 9 FOVs per n per condition. Data for G1 and G2 cells have passed Shapiro-Wilk test for Gaussian distribution. Statistical significance was determined using unpaired two-tailed t-test; G1 cells (-Dox d3 vs +Dox d3): P = 0.0036. G2 cells (-Dox d3 vs +Dox d3): P = 0.0034. Mann-Whitney test was used for Early S (-Dox d3 vs +Dox d3): P = 0.2084, ns.

**Supplementary Figure 2. MCF10A cells transition from E to early E/M when treated with TGFβ1.**

**A.** Cell transition from E to early E/M upon TGFβ1 treatment. Created in BioRender.

**B.** Fluorescent gelatin layer (left) and corresponding phase-contrast images of MCF10A cells, with or without TGFβ1 treatment, and with lovastatin-driven synchronization to G1 phase. Scale bar, 50 µm.

**C.** Quantification of matrix degradation area per cell corresponding to (B). Dotted lines represent quartiles; the dashed line represents the median. Statistical significance was determined using the unpaired Mann–Whitney test. ***P < 0.001; ****P < 0.0001. N=2, n=1, 9-10 FOVs per n per condition.

**D.** Western blot analysis of E and M marker expression in MCF10A cells before and after TGFβ1 treatment.

**Supplementary Figure 3. Differential expression of matrix remodeling genes across cell cycle phases in early- and late E/M states.**

**A.** Heatmap showing the relative expression levels of statistically significant matrix remodeling and invadopodia- related genes in early E/M cells, G1 versus G2 cell cycle phases.

**B.** Heatmap showing the relative expression levels of statistically significant matrix remodeling and invadopodia- related genes in late E/M cells, G1 versus G2 cell cycle phases. The color scale represents Z-score normalized expression values across conditions.

**Supplementary Figure 4. Differential expression of matrix remodeling genes between early- and late E/M cells in the same cell cycle phases.**

**A.** Heatmap showing the relative expression levels of statistically significant matrix remodeling and invadopodia- related genes in G1 phase: early- vs late E/M cells.

**B.** Heatmap showing the relative expression levels of statistically significant matrix remodeling and invadopodia- related genes in G2 phase: early- vs late E/M cells. The color scale represents Z-score normalized expression values across conditions.

**Supplementary Figure 5. High FILIP1L expression is associated with increased expression of ECM-degrading proteases.**

**A-F**. Violin plots showing lo G2 mRNA expression of MMPs (**a**-**b**), invadopodia components (**c**), ADAMs (**d**), ADAM, ADAMEC and ADAMTS family proteases (**e**-**f**) in breast cancer patient samples stratified by FILIP1L expression (low: Group A, blue; high: Group B, pink). Data was retrieved from publicly available breast cancer TCGA PanCancer Atlas dataset. Dotted lines represent quartiles; the dashed line represents the median. Significance was determined using the unpaired Mann-Whitney test. P values: P>0.05 (ns), *P*□<□0.05 (\**), P*U*<*U*0.01 (**), P*U*<*U*0.001 (\*\*\**), *P*□<□0.0001 (****).

**Supplementary Figure 6. FILIP1L colocalizes with Tks5 and cortactin in MDA-MB-231 cells**

**A.** In MDA-MB-231 cells labeled by DAPI (blue), Tks5 (green) and FILIP1L (red), white arrows point to invadopodia. Line (magenta) was drawn through one of the invadopodia punctae. Right panel: Profile of FILIP1L (red) and Tks5 (green) intensities along the magenta line Scale bar: 20 µm.

**B.** In MDA-MB-231 cells labeled by DAPI (blue), cortactin (green) and FILIP1L (red), white arrows point to invadopodia. Line (magenta) was drawn through one of the invadopodia punctae. Right panel: Profile of FILIP1L (red) and cortactin (green) intensities along the magenta line. Scale bar: 20 µm.

## References

1. Survival Rates for Breast Cancer [Internet]. [cited 2025 Nov 5]. https://www.cancer.org/cancer/types/breast-cancer/understanding-a-breast-cancer-diagnosis/breast-cancer-survival-rates.html. Accessed 5 Nov 2025

2. DeSantis CE, Ma J, Gaudet MM, Newman LA, Miller KD, Goding Sauer A, et al. Breast cancer statistics, 2019. CA Cancer J Clin. 2019;69:438–51. 10.3322/caac.21583

3. Ording AG, Heide-Jørgensen U, Christiansen CF, Nørgaard M, Acquavella J, Sørensen HT. Site of metastasis and breast cancer mortality: a Danish nationwide registry-based cohort study. Clin Exp Metastasis. 2017;34:93–101. 10.1007/s10585-016-9824-8

4. Cummings MC, Simpson PT, Reid LE, Jayanthan J, Skerman J, Song S, et al. Metastatic progression of breast cancer: insights from 50 years of autopsies. J Pathol. 2014;232:23–31. 10.1002/path.4288

5. Gligorijevic B, Bergman A, Condeelis J. Multiparametric Classification Links Tumor Microenvironments with Tumor Cell Phenotype. PLOS Biol. Public Library of Science; 2014;12:e1001995. 10.1371/journal.pbio.1001995

6. Williams KC, Cepeda MA, Javed S, Searle K, Parkins KM, Makela AV, et al. Invadopodia are chemosensing protrusions that guide cancer cell extravasation to promote brain tropism in metastasis. Oncogene. 2019;38:3598–615. 10.1038/s41388-018-0667-4

7. Gligorijevic B, Wyckoff J, Yamaguchi H, Wang Y, Roussos ET, Condeelis J. N-WASP-mediated invadopodium formation is involved in intravasation and lung metastasis of mammary tumors. J Cell Sci. 2012;125:724–34. 10.1242/jcs.092726

8. Leong HS, Robertson AE, Stoletov K, Leith SJ, Chin CA, Chien AE, et al. Invadopodia Are Required for Cancer Cell Extravasation and Are a Therapeutic Target for Metastasis. Cell Rep. Cell Press; 2014;8:1558–70. 10.1016/J.CELREP.2014.07.050

9. Kalluri R, Weinberg RA. The basics of epithelial-mesenchymal transition. J Clin Invest. 2009;119:1420–8. 10.1172/JCI39104

10. Debnath P, Huirem RS, Dutta P, Palchaudhuri S. Epithelial–mesenchymal transition and its transcription factors. Biosci Rep. 2021;42:BSR20211754. 10.1042/BSR20211754

11. Wang J, Wei Q, Wang X, Tang S, Liu H, Zhang F, et al. Transition to resistance: An unexpected role of the EMT in cancer chemoresistance. Genes Dis. 2016;3:3–6. 10.1016/j.gendis.2016.01.002

12. Zhang B, Zhao R, Wang Q, Zhang Y-J, Yang L, Yuan Z-J, et al. An EMT-Related Gene Signature to Predict the Prognosis of Triple-Negative Breast Cancer. Adv Ther. 2023;40:4339–57. 10.1007/s12325-023-02577-z

13. Ribatti D, Tamma R, Annese T. Epithelial-Mesenchymal Transition in Cancer: A Historical Overview. Transl Oncol. 2020;13:100773. 10.1016/j.tranon.2020.100773

14. Debaugnies M, Rodríguez-Acebes S, Blondeau J, Parent M-A, Zocco M, Song Y, et al. RHOJ controls EMT-associated resistance to chemotherapy. Nature. Nature Publishing Group; 2023;616:168–75. 10.1038/s41586-023-05838-7

15. Sundararajan V, Gengenbacher N, Stemmler MP, Kleemann JA, Brabletz T, Brabletz S. The ZEB1/miR-200c feedback loop regulates invasion via actin interacting proteins MYLK and TKS5. Oncotarget. 2015;6:27083–96.

16. Perrin L, Belova E, Bayarmagnai B, Tüzel E, Gligorijevic B. Invadopodia enable cooperative invasion and metastasis of breast cancer cells. Commun Biol. Nature Publishing Group; 2022;5:1–14. 10.1038/s42003-022-03642-z

17. Hanrahan K, O’Neill A, Prencipe M, Bugler J, Murphy L, Fabre A, et al. The role of epithelial–mesenchymal transition drivers ZEB1 and ZEB2 in mediating docetaxel-resistant prostate cancer. Mol Oncol. 2017;11:251–65. 10.1002/1878-0261.12030

18. Weiss MB, Abel EV, Mayberry MM, Basile KJ, Berger AC, Aplin AE. TWIST1 Is an ERK1/2 Effector That Promotes Invasion and Regulates MMP-1 Expression in Human Melanoma Cells. Cancer Res. 2012;72:6382–92. 10.1158/0008-5472.CAN-12-1033

19. Lin C-Y, Tsai P-H, Kandaswami CC, Lee P-P, Huang C-J, Hwang J-J, et al. Matrix metalloproteinase-9 cooperates with transcription factor Snail to induce epithelial–mesenchymal transition. Cancer Sci. 2011;102:815–27. 10.1111/j.1349-7006.2011.01861.x

20. Sinha D, Saha P, Samanta A, Bishayee A. Emerging Concepts of Hybrid Epithelial-to-Mesenchymal Transition in Cancer Progression. Biomolecules. 2020;10:1561. 10.3390/biom10111561

21. Pastushenko I, Blanpain C. EMT Transition States during Tumor Progression and Metastasis. Trends Cell Biol. Elsevier; 2019;29:212–26. 10.1016/j.tcb.2018.12.001

22. Pastushenko I, Mauri F, Song Y, de Cock F, Meeusen B, Swedlund B, et al. Fat1 deletion promotes hybrid EMT state, tumour stemness and metastasis. Nature. Nature Publishing Group; 2021;589:448–55. 10.1038/s41586-020-03046-1

23. Perrin L, Gligorijevic B. Proteolytic and Mechanical Remodeling of the Extracellular Matrix by Invadopodia in Cancer. Phys Biol. 2022;20:10.1088/1478-3975/aca0d8. https://doi.org/10.1088/1478-3975/aca0d8

24. Pourfarhangi KE, Bergman A, Gligorijevic B. ECM Cross-Linking Regulates Invadopodia Dynamics. Biophys J. 2018;114:1455–66. 10.1016/j.bpj.2018.01.027

25. Bayarmagnai B, Perrin L, Pourfarhangi KE, Graña X, Tüzel E, Gligorijevic B. Invadopodia-mediated ECM degradation is enhanced in the G1 phase of the cell cycle. J Cell Sci [Internet]. Company of Biologists Ltd; 2019 [cited 2023 Aug 21];132. 10.1242/JCS.227116

26. Wolf K, te Lindert M, Krause M, Alexander S, te Riet J, Willis AL, et al. Physical limits of cell migration: Control by ECM space and nuclear deformation and tuning by proteolysis and traction force. J Cell Biol. 2013;201:1069–84. 10.1083/jcb.201210152

27. Perrin L, Tucker T, Gligorijevic B. Time-Resolved Fluorescence Imaging and Analysis of Cancer Cell Invasion in the 3D Spheroid Model. J Vis Exp JoVE. 2021;e61902. 10.3791/61902

28. Gong Z, Wisdom KM, McEvoy E, Chang J, Adebowale K, Price CC, et al. Recursive feedback between matrix dissipation and chemo-mechanical signaling drives oscillatory growth of cancer cell invadopodia. Cell Rep. 2021;35:109047. 10.1016/j.celrep.2021.109047

29. Gould CM, Courtneidge SA. Regulation of invadopodia by the tumor microenvironment. Cell Adhes Migr. 2014;8:226–35. 10.4161/cam.28346

30. Lehn S, Tobin NP, Berglund P, Nilsson K, Sims AH, Jirström K, et al. Down-Regulation of the Oncogene Cyclin D1 Increases Migratory Capacity in Breast Cancer and Is Linked to Unfavorable Prognostic Features. Am J Pathol. American Society for Investigative Pathology; 2010;177:2886. 10.2353/AJPATH.2010.100303

31. Walmod PS, Hartmann-Petersen R, Prag S, Lepekhin EL, Röpke C, Berezin V, et al. Cell-cycle-dependent regulation of cell motility and determination of the role of Rac1. Exp Cell Res. 2004;295:407–20. 10.1016/j.yexcr.2004.01.011

32. Zhang J, Goliwas KF, Wang W, Taufalele PV, Bordeleau F, Reinhart-King CA. Energetic regulation of coordinated leader–follower dynamics during collective invasion of breast cancer cells. Proc Natl Acad Sci. Proceedings of the National Academy of Sciences; 2019;116:7867–72. 10.1073/pnas.1809964116

33. Moshfegh Y, Bravo-Cordero JJ, Miskolci V, Condeelis J, Hodgson L. A Trio-Rac1-PAK1 signaling axis drives invadopodia disassembly. Nat Cell Biol. 2014;16:574–86. 10.1038/ncb2972

34. Brown KA, Aakre ME, Gorska AE, Price JO, Eltom SE, Pietenpol JA, et al. Induction by transforming growth factor-β1 of epithelial to mesenchymal transition is a rare event in vitro. Breast Cancer Res. 2004;6:R215–31. 10.1186/bcr778

35. Xu J, Lamouille S, Derynck R. TGF-β-induced epithelial to mesenchymal transition. Cell Res. 2009;19:156–72. 10.1038/cr.2009.5

36. Guo W, Keckesova Z, Donaher JL, Shibue T, Tischler V, Reinhardt F, et al. Slug and Sox9 cooperatively determine the mammary stem cell state. Cell. 2012;148:1015–28. 10.1016/j.cell.2012.02.008

37. Phillips S, Kuperwasser C. SLUG: Critical regulator of epithelial cell identity in breast development and cancer. Cell Adhes Migr. 2014;8:578–87. 10.4161/19336918.2014.972740

38. Elloul S, Bukholt Elstrand M, Nesland JM, Tropé CG, Kvalheim G, Goldberg I, et al. Snail, Slug, and Smad-interacting protein 1 as novel parameters of disease aggressiveness in metastatic ovarian and breast carcinoma. Cancer. 2005;103:1631–43. 10.1002/cncr.20946

39. Côme C, Magnino F, Bibeau F, De Santa Barbara P, Becker KF, Theillet C, et al. Snail and Slug Play Distinct Roles during Breast Carcinoma Progression. Clin Cancer Res. 2006;12:5395–402. 10.1158/1078-0432.CCR-06-0478

40. Pignatelli J, Tumbarello DA, Schmidt RP, Turner CE. Hic-5 promotes invadopodia formation and invasion during TGF-β–induced epithelial–mesenchymal transition. J Cell Biol. 2012;197:421–37. 10.1083/jcb.201108143

41. Wolf K, te Lindert M, Krause M, Alexander S, te Riet J, Willis AL, et al. Physical limits of cell migration: Control by ECM space and nuclear deformation and tuning by proteolysis and traction force. J Cell Biol. 2013;201:1069–84. 10.1083/jcb.201210152

42. Juang Y-L, Jeng Y-M, Chen C-L, Lien H-C. PRRX2 as a novel TGF-β-induced factor enhances invasion and migration in mammary epithelial cell and correlates with poor prognosis in breast cancer. Mol Carcinog. 2016;55:2247–59. 10.1002/mc.22465

43. Wendt MK, Smith JA, Schiemann WP. Transforming Growth Factor-β-Induced Epithelial-Mesenchymal Transition Facilitates Epidermal Growth Factor-Dependent Breast Cancer Progression. Oncogene. 2010;29:6485–98. 10.1038/onc.2010.377

44. Bugaeva O, Maliniemi P, Prestvik WS, Leivo E, Kluger N, SALAVA A, et al. Tumour Suppressor Neuron Navigator 3 and Matrix Metalloproteinase 14 are Co-expressed in Most Melanomas but Downregulated in Thick Tumours. Acta Derm Venereol. 2023;103:298. 10.2340/actadv.v103.298

45. Uboveja A, Satija YK, Siraj F, Sharma I, Saluja D. p73 – NAV3 axis plays a critical role in suppression of colon cancer metastasis. Oncogenesis. Nature Publishing Group; 2020;9:1–15. 10.1038/s41389-020-0193-4

46. Zhang H, Pan Y, Cheung M, Cao M, Yu C, Chen L, et al. LAMB3 mediates apoptotic, proliferative, invasive, and metastatic behaviors in pancreatic cancer by regulating the PI3K/Akt signaling pathway. Cell Death Dis. 2019;10:230. 10.1038/s41419-019-1320-z

47. Medici D, Nawshad A. Type I collagen promotes epithelial-mesenchymal transition through ILK-dependent activation of NF-κB and LEF-1. Matrix Biol J Int Soc Matrix Biol. 2010;29:161–5. 10.1016/j.matbio.2009.12.003

48. Jung S-N, Lim HS, Liu L, Chang JW, Lim YC, Rha KS, et al. LAMB3 mediates metastatic tumor behavior in papillary thyroid cancer by regulating c-MET/Akt signals. Sci Rep. Nature Publishing Group; 2018;8:2718. 10.1038/s41598-018-21216-0

49. Bonnans C, Chou J, Werb Z. Remodelling the extracellular matrix in development and disease. Nat Rev Mol Cell Biol. 2014;15:786–801. 10.1038/nrm3904

50. Wang LW, Nandadasa S, Annis DS, Dubail J, Mosher DF, Willard BB, et al. A disintegrin-like and metalloproteinase domain with thrombospondin type 1 motif 9 (ADAMTS9) regulates fibronectin fibrillogenesis and turnover. J Biol Chem. Elsevier; 2019;294:9924–36. 10.1074/jbc.RA118.006479

51. Arora S, Scott AM, Janes PW. ADAM Proteases in Cancer: Biological Roles, Therapeutic Challenges, and Emerging Opportunities. Cancers. 2025;17:1703. 10.3390/cancers17101703

52. Huang Y, Hong W, Wei X. The molecular mechanisms and therapeutic strategies of EMT in tumor progression and metastasis. J Hematol OncolJ Hematol Oncol. 2022;15:129. 10.1186/s13045-022-01347-8

53. Gilles C, Newgreen DF, Sato H, Thompson EW. Matrix Metalloproteases and Epithelial-to-Mesenchymal Transition: Implications for Carcinoma Metastasis. Madame Curie Biosci Database Internet [Internet]. Landes Bioscience; 2013 [cited 2025 May 2]. https://www.ncbi.nlm.nih.gov/books/NBK6387/. Accessed 2 May 2025

54. Peixoto P, Etcheverry A, Aubry M, Missey A, Lachat C, Perrard J, et al. EMT is associated with an epigenetic signature of ECM remodeling genes. Cell Death Dis. Nature Publishing Group; 2019;10:1–17. 10.1038/s41419-019-1397-4

55. Shen C-J, Kuo Y-L, Chen C-C, Chen M-J, Cheng Y-M. MMP1 expression is activated by Slug and enhances multi-drug resistance (MDR) in breast cancer. PLOS ONE. Public Library of Science; 2017;12:e0174487. 10.1371/journal.pone.0174487

56. Wendt MK, Allington TM, Schiemann WP. Mechanisms of the epithelial-mesenchymal transition by TGF-beta. Future Oncol Lond Engl. 2009;5:1145–68. 10.2217/fon.09.90

57. Henretta S, Bockley K, Odell J, Lammerding J. Filamin A interacting protein 1-like (FILIP1L) has mitochondrial localization. MicroPublication Biol. 2025:10.17912/micropub.biology.001572. https://doi.org/10.17912/micropub.biology.001572

58. Kwon M, Lee SJ, Wang Y, Rybak Y, Luna A, Reddy S, et al. Filamin A interacting protein 1-like inhibits WNT signaling and MMP expression to suppress cancer cell invasion and metastasis. Int J Cancer J Int Cancer. 2014;135:48–60. 10.1002/ijc.28662

59. Kwon M, Kim J-H, Rybak Y, Luna A, Choi CH, Chung J-Y, et al. Reduced expression of FILIP1L, a novel WNT pathway inhibitor, is associated with poor survival, progression and chemoresistance in ovarian cancer. Oncotarget. Impact Journals; 2016;7:77052–70. 10.18632/oncotarget.12784

60. Nathanson SD, Detmar M, Padera TP, Yates LR, Welch DR, Beadnell TC, et al. Mechanisms of breast cancer metastasis. Clin Exp Metastasis. 2022;39:117–37. 10.1007/s10585-021-10090-2

61. Lengyel E. Ovarian Cancer Development and Metastasis. Am J Pathol. 2010;177:1053–64. 10.2353/ajpath.2010.100105

62. Richner M, Victor MB, Liu Y, Abernathy D, Yoo AS. MicroRNA-based conversion of human fibroblasts into striatal medium spiny neurons. Nat Protoc. Nature Publishing Group; 2015;10:1543–55. 10.1038/nprot.2015.102

63. Guo W, Keckesova Z, Donaher JL, Shibue T, Tischler V, Reinhardt F, et al. Slug and Sox9 cooperatively determine the mammary stem cell state. Cell. 2012;148:1015–28. 10.1016/j.cell.2012.02.008

64. Sharma VP, Entenberg D, Condeelis J. High-Resolution Live-Cell Imaging and Time-Lapse Microscopy of Invadopodium Dynamics and Tracking Analysis. In: Coutts AS, editor. Adhes Protein Protoc [Internet]. Totowa, NJ: Humana Press; 2013 [cited 2025 June 29]. p. 343–57. 10.1007/978-1-62703-538-5_21

65. Lloyd-Lewis B, Davis FM, Harris OB, Hitchcock JR, Lourenco FC, Pasche M, et al. Imaging the mammary gland and mammary tumours in 3D: optical tissue clearing and immunofluorescence methods. Breast Cancer Res. 2016;18:127. 10.1186/s13058-016-0754-9

