## Supplementary Tables for "EMT and cell cycle control invadopodia and metastasis in breast cancer via Filip1L"

**Supplementary Table 1. DESeq2 design setup**

| Samples | Treatment | Cell cycle phase | Treatment_color | Invasiveness |
|---|---|---|---|---|
| Early E/M G2 #1 | -TGFβ-1 (Ctl) | G2 (Green) | Ctl_G | Invasive |
| Early E/M G1 #1 | | G1 (Red) | Ctl_R | Noninvasive |
| Early E/M G2 #2 | | G2 (G) | Ctl_G | Invasive |
| Early E/M G1 #2 | | G1 (R) | Ctl_R | Noninvasive |
| Early E/M G2 #3 | | G2 (G) | Ctl_G | Invasive |
| Early E/M G1 #3 | | G1 (R) | Ctl_R | Noninvasive |
| Early E/M G2 #4 | | G2 (G) | Ctl_G | Invasive |
| Early E/M G1 #4 | | G1 (R) | Ctl_R | Noninvasive |
| Late E/M G2 #1 | +TGFβ-1 | G2 (G) | Growth_G | Noninvasive |
| Late E/M G1 #1 | | G1 (R) | Growth_R | Invasive |
| Late E/M G2 #2 | | G2 (G) | Growth_G | Noninvasive |
| Late E/M G1 #2 | | G1 (R) | Growth_R | Invasive |
| Late E/M G2 #3 | | G2 (G) | Growth_G | Noninvasive |
| Late E/M G1 #3 | | G1 (R) | Growth_R | Invasive |
| Late E/M G2 #4 | | G2 (G) | Growth_G | Noninvasive |
| Late E/M G1 #4 | | G1 (R) | Growth_R | Invasive |

**Supplementary Table 2. Result of Multivariate DESeq2 analysis**

| Gene ID | baseMean | lo G2 FoldChange | p-value | padj | Symbol |
|---|---|---|---|---|---|
| ENSMUSG00000020897 | 2090.73 | -0.304 | 5.20E-08 | 0.0013 | Aurkb |
| ENSMUSG00000043336 | 1743.81 | 0.278 | 1.47E-07 | 0.0018 | FILIP1L |

**Supplementary Table 3. Materials for immunofluorescence**

| Antibody and dyes | Cat. no | Dilution |
|---|---|---|
| Anti-Tks5 | Millipore, MABT336 | 1:200 |
| Anti-E-cadherin | Thermo Fisher Scientific, 131900 | 1:200 |
| Anti-FILIP1L | Abcam, ab151331 | 1:200 |
| Vimentin | Abcam, ab92547 | 1:200 |
| Phalloidin | Life Technologies<br><br>A22283, A22284 | 1:250 |

**Supplementary Table 4. Materials for Western blots**

| I Antibody | Cat. # | Host | I Antibody Dilution | II Antibody Dilution | µg of protein/lane | Chemiluminescent reagent |
|---|---|---|---|---|---|---|
| Vimentin | Abcam, ab92547 | Rabbit | 1:1000 | 1:4000 | 10 | WesternBright |
| FILIP1L | Abcam, ab151331 | Rabbit | 1:1000 | 1:4000 | 10 | WesternBright |
| FILIP1L Thermo | PA5-60251 | Rabbit | 1:1000 | 1:5000 | 25 | WesternBright |
| Slug (C19G7) | Cell Signaling Technology 9585T | Rabbit | 1:1000 | 1:5000 | 20 | WesternBright |
| E-cadherin BD | BD Biosciences, 610181 | Mouse | 1:1000 | 1:4000 | 10 | WesternBright |
| β-actin Loading Control Monoclonal Antibody (BA3R) | Thermo Scientific MA5-15739 | Mouse | 1:1000 | 1:5000 | 10 | WesternBright |
| ZO1 | Thermo Fisher, 21773-1-AP | Rabbit | 1:1000 | 1:5000 | 10 | WesternBright |
| Tks5 | MABT336 | Mouse | 1:100 | 1:5000 | 10 | Supersignal |
| Twist1 | Sigma, T6451-25UL | Rabbit | 1:1000 | 1:4000 | 10 | Super signal |
