## Supplementary Figures for "EMT and cell cycle control invadopodia and metastasis in breast cancer via Filip1L"

**Supplementary Figure 1.**

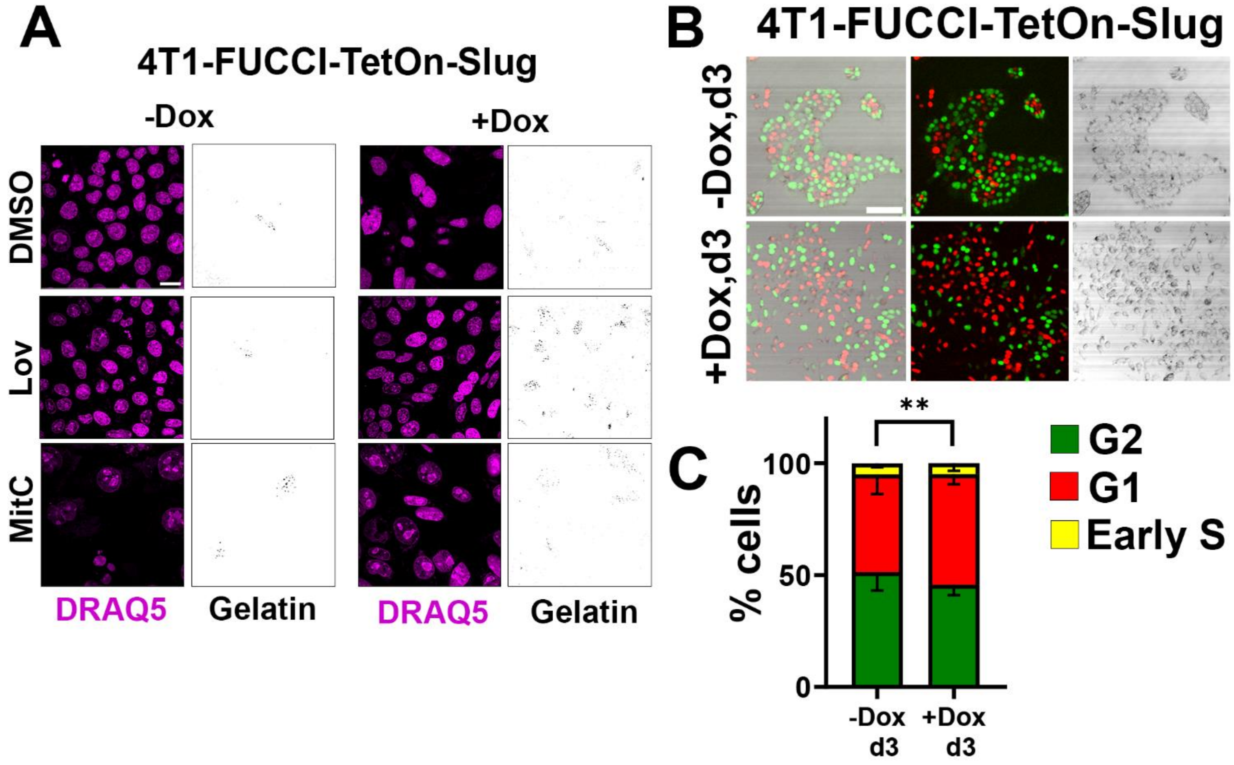

Supplementary Figure 2.

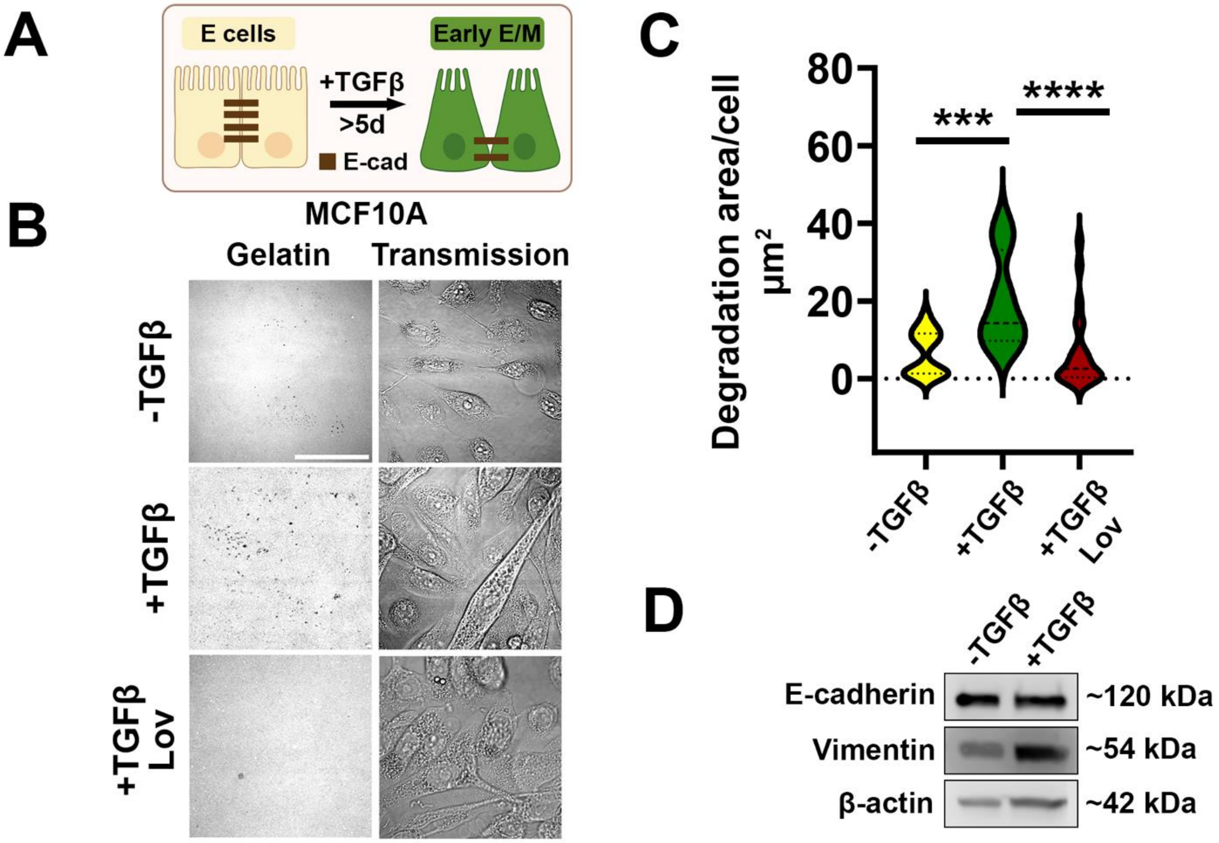

**Supplementary Figure 3.**

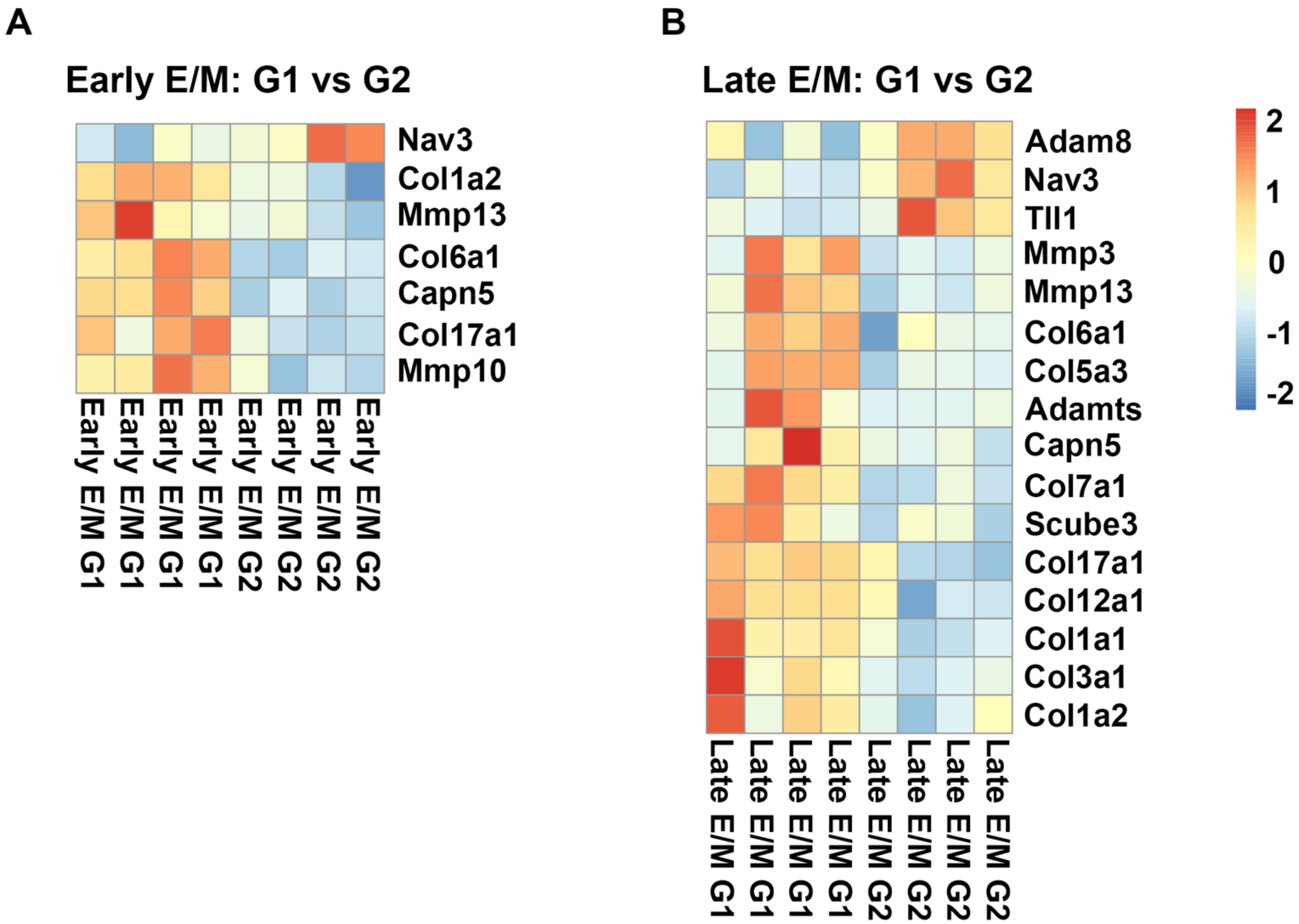

Supplementary Figure 4.

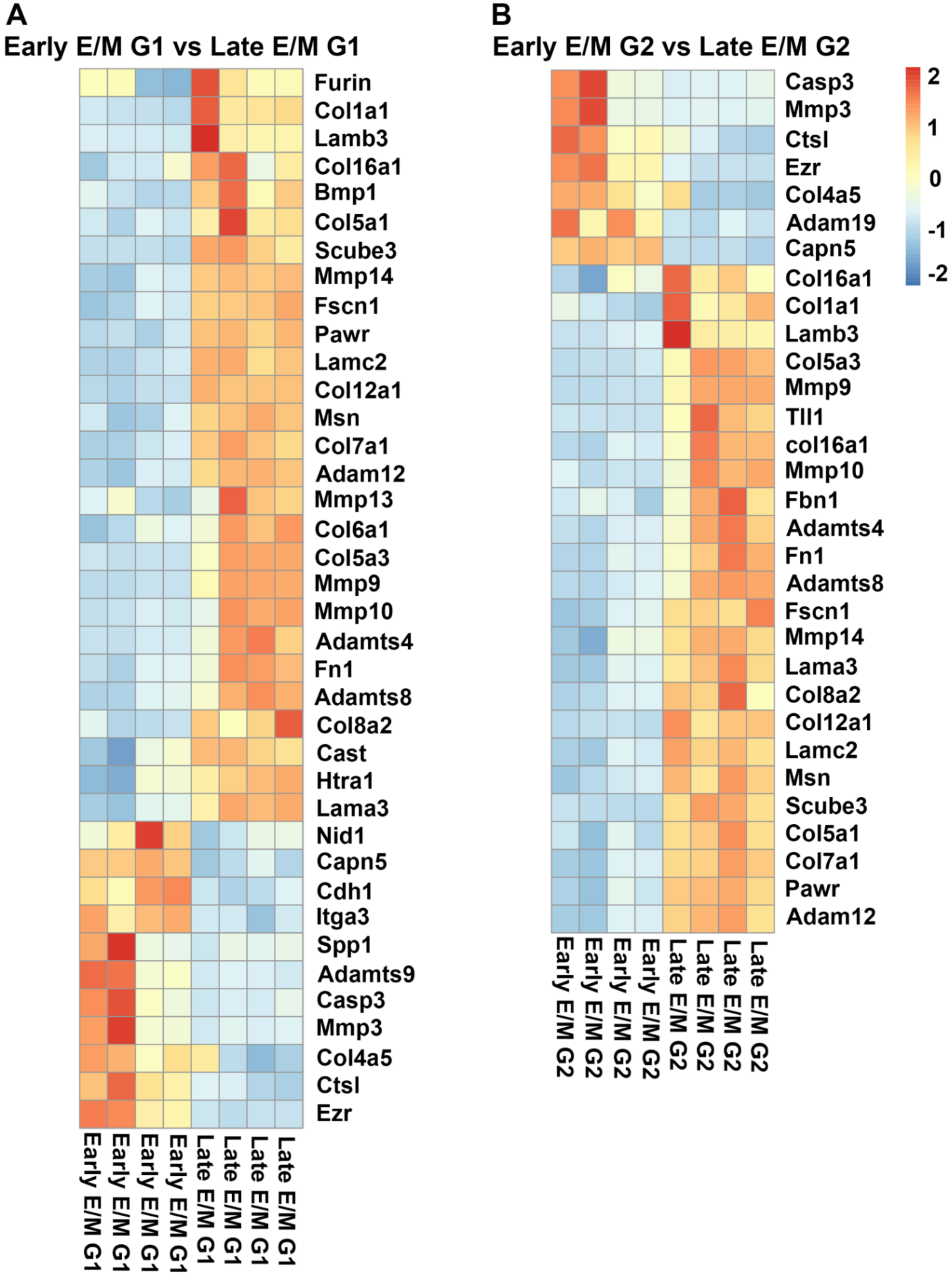

**Supplementary Figure 5.**

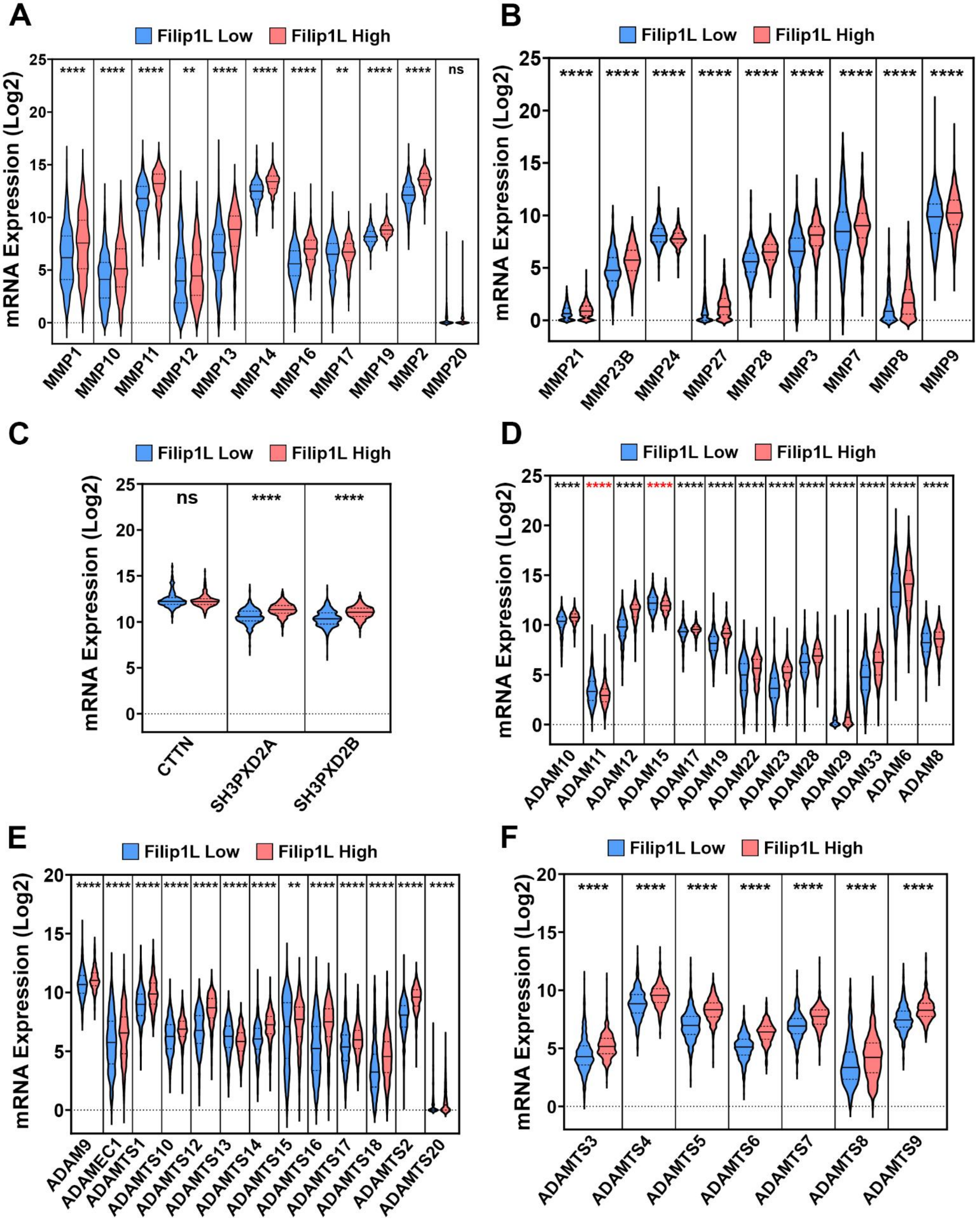

**Supplementary Figure 6.**

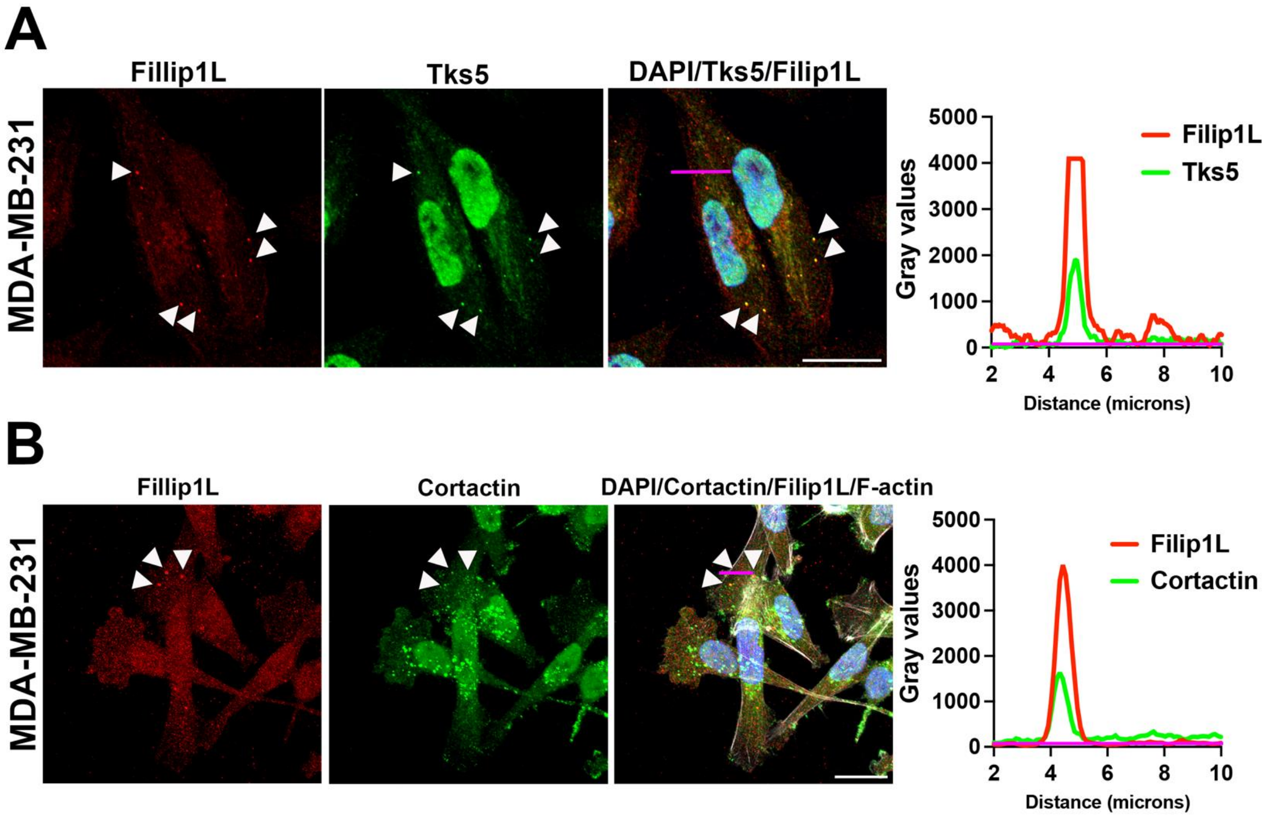
